# Input-and cell-type-specific developmental alterations to thalamic synapses in a Dravet syndrome mouse model

**DOI:** 10.64898/2026.02.12.705567

**Authors:** Mona Safari, Rutvi Desai, Harbal Rai, Tyler J. Roberts, Rabeya Khondaker, Julia Smith, Sharon A. Swanger

**Author notes:** Corresponding Author: Sharon A. Swanger, Fralin Biomedical Research Institute at VTC Center for Neurobiology Research, 2 Riverside Circle, Roanoke, VA 24016.

## Abstract

Dravet syndrome is an epileptic encephalopathy most often caused by loss-of-function mutations in the *SCN1A* gene, leading to haploinsufficiency of the voltage-gated sodium channel Na_V_1.1. Seizures begin during infancy and generally wane throughout childhood, but behavioral symptoms, such as intellectual disability, motor impairments, and autistic features, remain through adulthood. Seizures primarily stem from inhibitory neuron hypo-excitability in the cortex, hippocampus, and thalamus, but circuit abnormalities underlying persistent behavioral symptoms are poorly understood. Prior work showed synapse dysfunction in thalamocortical neurons in four-week-old DS mice. To understand when synaptic deficits develop and whether they could contribute to persistent thalamic dysfunction, we investigated synapse function in the ventral posterolateral (VPL) and ventral posteromedial (VPM) thalamus prior to seizure onset (P13-P17), after the period of highest seizure burden (P28-P32), and in adulthood (P58-P63). Recordings of VPL and VPM synaptic activity showed excitatory input to the VPL was significantly reduced after seizure onset and this reduction persisted through adulthood, while VPM excitatory input was unaffected. We further showed a selective reduction in the function and number of excitatory sensory synapses in the VPL, with no change to cortical synapses. VPL and VPM neurons both showed inhibitory synapse dysfunction at four weeks, which persisted into adult DS mice only in VPL neurons. These results revealed persistent input- and cell-type-specific alterations to thalamic synapses that develop after seizure onset and are maintained into adulthood, suggesting that synaptic deficits could contribute to ongoing circuit dysfunction in DS.

## INTRODUCTION

Dravet syndrome (DS) is a severe form of myoclonic epilepsy that is most often caused by mutations in the *SCN1A* gene, which encodes the voltage-gated sodium channel Na_V_1.1 (Hurst 1990; Wu et al. 2015). DS has an incidence of approximately 1 in 80,000 people in the U.S., and the patient mortality rate is 15-20% due to sudden unexpected death in epilepsy (SUDEP) resulting from prolonged seizures (Cooper et al. 2016; Dravet 2011; Skluzacek et al. 2011). Na_V_1.1 facilitates action potential generation and propagation via expression at the axon initial segment and nodes of Ranvier (Duflocq et al. 2008; Favero et al. 2018; Hedrich et al. 2014). The *Scn1a* gene is highly expressed in GABAergic neurons in the cortex, hippocampus, and reticular thalamus (nRT) all of which display significant reduction in excitability in DS mouse models (Cheah et al. 2012; Kalume et al. 2015; Ogiwara et al. 2007; Yu et al. 2006a). The resulting disinhibition leads to circuit hyperexcitability beginning in the third post-natal week contributing to spontaneous convulsive seizures, heat-induced seizures, and spike-and-wave discharges (Cheah et al. 2012; Yu et al. 2006a; Oakley et al. 2009). This phenotype recapitulates the human disease involving myoclonic jerks, tonic-clonic seizures, febrile seizures, and absence seizures with a median onset of 6 months (Dravet, 2011a; Jansson et al., 2020; Kalume et al., 2015; Li et al., 2011).

Seizures lessen throughout adolescence into adulthood in patients with DS, but debilitating behavioral afflictions continue including motor impairments, intellectual disability, sleep impairment, and autistic behaviors. While much has been learned about aberrant spike generation underlying seizures in DS, far less is known about the enduring mechanisms contributing to behavioral impairments throughout the lifespan. Given that the highest seizure burden occurs during postnatal development, activity-dependent synapse formation is likely disrupted in DS, which could result in miswiring of brain circuits and perpetual brain dysfunction. Notably, cortical parvalbumin interneuron spike generation normalizes by adulthood in DS mouse models, but parvalbumin GABAergic synaptic transmission remains disrupted (Favero et al., 2018; Kaneko et al., 2022). Impaired action potential propagation in axons contributes to this sustained disruption in synapse function, but other studies have suggested that altered synaptic transmission or development may contribute to disrupted inhibition in DS (Uchino et al. 2021; Chancey and Howard 2022; Layer et al. 2023; Studtmann et al. 2022). Recent work in a human induced pluripotent cell line from a patient with DS demonstrated that network dysfunction involves reduced excitatory synapse strength, and computational modeling further supports the likelihood of homeostatic downscaling of excitatory synapses in DS models (Doorn et al., 2023). On the other hand, enhanced excitatory input to dentate gyrus granule cells contributes to persistent excessive excitation in hippocampal circuits (Mattis et al., 2022). While several reports indicate that altered synaptic transmission may contribute to circuit dysfunction in DS, the direction, magnitude, and duration of these changes as well as the specific synaptic connections affected remain largely unresolved.

In the thalamus, Na_V_1.1 is expressed in glutamatergic thalamocortical (TC) neurons in the ventral posterolateral (VPL) and ventral posteromedial (VPM) thalamus as well as the GABAergic nRT (Studtmann et al., 2022). In addition to the profound alterations in inhibitory nRT neuron excitability (Hedrich et al., 2014; Ritter-Makinson et al., 2019; Studtmann et al., 2022), TC neurons exhibit cell-type- and synapse-specific alterations that may further disrupt thalamocortical network function in DS (Studtmann et al., 2022). VPL and VPM neurons receive glutamatergic input from corticothalamic neurons, glutamatergic sensory input from the spinal cord and brainstem, respectively, and GABAergic input from the nRT. In a DS mouse model, VPL neurons showed reduced excitability as well as reduced glutamatergic sensory input, while VPM neurons exhibited increased excitability and no apparent changes in synaptic input (Studtmann et al., 2022). Notably, CT input to the nRT was reduced, while CT input to VPL and VPM was unchanged. This cell-type- and synapse-specific pathophysiology within the somatosensory CT circuitry indicates that hyperexcitable circuits do not undergo a global homeostatic scaling of synaptic transmission in DS. However, the underlying mechanisms and timeline of cell-type- and input-specific dysfunction remain unknown.

To better understand how synaptic transmission may be disrupted within the somatosensory thalamus, we investigated if altered synapse development occurs before seizure onset, potentially contributing to circuit dysfunction underlying seizures, or after the period of highest seizure burden indicating that it might be a consequence of network hyperexcitability. We further examined if thalamic synapse dysfunction was maintained into adulthood. In this study, we utilized electrophysiology and high-resolution imaging to selectively quantify glutamatergic CT and sensory inputs as well as GABAergic inputs to VPL and VPM neurons. This work uncovered that input-and cell-type-specific synapse dysfunction in VPL neurons developed after seizure onset and some, but not all, of the altered synaptic connections showed enduring changes in adult DS mice.

## METHODS

### Mouse model

Mouse studies were used according to the protocols approved by the Institutional Animal Care and Use Committee at Virginia Polytechnic Institute and State University and in accordance with the National Institutes of Health guidelines. We crossed heterozygous 129S6/SvEvTac-Scn1a^tm1Kea/WT^ mice with C57BL/6J mice. Scn1a^+/+^ and Scn1a^+/-^littermates of both sexes were used at the age of P13-P17, P28-P32, and P58-P62. Transnetyx performed mouse genotyping by using real-time PCR.

### Electrophysiology

For recording synaptic activity, mice were euthanized by isoflurane overdose and then perfused with ice-cold sucrose-based artificial cerebrospinal fluid (aCSF) containing (in mM) 230 sucrose, 24 NaHCO3, 10 glucose, 3 KCl, 10 MgSO4, 1.25 NaH2PO4, and 0.5 CaCl2 saturated with 95% O2 / 5% CO2. The brain was removed and sliced (300 µm) with the vibratome (Leica VT1200S). Slices were cut horizontally in the oxygenated cold sucrose aCSF. Slices were incubated in the NaCl-based aCSF containing (in mM) 130 NaCl, 24 NaHCO3, 10 glucose, 3 KCl, 4 MgSO4, 1.25 NaH2PO4, and 0.5 CaCl2 saturated with 95% O2 / 5% CO2 at room temperature.

One recording was performed per slice and no more than two cells per mouse were included in a single dataset. Recordings were made using a Multiclamp 700B amplifier (Molecular Devices), sampled at 20 kHz (Digi-data 1550B, Molecular Devices), and low pass filtered at 10 kHz using Axon pClamp 11 software (Molecular Devices). The extracellular recording solution contained (in mM) 130 NaCl, 24 NaHCO3, 10 glucose, 3 KCl, 1 MgSO4, 1.25 NaH2PO4, and 2 CaCl2 saturated with 95% O2 / 5% CO2 and was maintained at 32 ◦C for all recordings. Whole-cell voltage-clamp recordings were made using borosilicate glass recording electrodes (4–5 MΩ) filled with (in mM) 120 CsMeSO3, 15 CsCl, 8 NaCl, 10 tetraethylammonium chloride, 10 HEPES, 1 EGTA, 3 Mg-ATP, 0.3 Na-GTP, 1.5 MgCl2, 1 QX 314, and 0.1% biocytin, pH 7.3. After a 10-minute equilibration period, sEPSCs and sIPSCs were recorded in two-minute epochs at a holding potential of -70 mV and 0 mV, respectively in the presence of the recording NaCl aCSF in VPL and VPM regions. For recording mEPSCs and mIPSCs, 1 μM TTX was added to the extracellular recording solution. Holding commands were adjusted for a 10-mV liquid junction potential during the recordings. Series resistance and cell capacitance were monitored throughout the experiment but were not compensated, and cells were excluded if either parameter changed > 20%.

### Electrophysiology data analysis

For sE(I)PSCs and mE(I)PSCs analysis, recordings of 3-6 minutes were analyzed by MiniAnalysis software (Synaptosoft) to determine the frequency and amplitude of the events. Data were digitally filtered at 1 kHz. Automated detection identified events with amplitude ≥7 pA (∼5 x RMS noise), and then events ≥5 pA were manually detected, and automated detection accuracy was assessed. The amplitude and frequency of the recordings averaged in 30-second bins in each recording. The decay time for all mEPSC events were measured by fitting 30 – 70% decay in MiniAnalysis and plotted as a frequency distribution for each cell independently. All distributions were multimodal (p < 0.05). Each distribution was interpolated using the Akima spline method (GraphPad Prism 9), and the first derivative of the spline was plotted to determine the x-intercept between the two modes, which is the decay time corresponding to the critical value used to separate the bimodal distribution two groups of mEPSC events. The events were separated into Type 1 (shorter, putative sensory events) and Type 2 (longer, putative cortical events) mEPSCs.

### Immunohistochemistry and imaging

For counting the number of glutamatergic synapses, mice were euthanized by isoflurane overdose, transcardially perfused with 4% paraformaldehyde (PFA), and post-fixed by 4% PFA for 24 hours. Brains were then transferred into 30% sucrose in 1× phosphate-buffered saline (PBS) for 48 h and frozen in OCT. Brains were cryosectioned (20 μm) and slide-mounted, and immunostained as previously described (Studtmann et al. 2022). Briefly, brain slices were treated with 0.8% sodium borohydride in 1× Tris-buffered saline (TBS) at room temperature, and then 0.01 M sodium citrate buffer at 100°C, pH 6.0 for 10 min. Permeabilized mounted brains with 0.5% Triton X-100 in 1×TBS, and blocked with FAB fragments and 10% normal donkey serum (NDS). Sections were incubated with antibodies against the following proteins overnight at 4°C: VGLUT1 (1:200, Synaptic Systems 135–511, RRID AB_887879), VGLUT2 (1:400, Synaptic Systems 135–402, AB_2187539), Homer (1:200; Synaptic Systems 160-003, RRID AB_887730; 1:200, Synaptic Systems 160–025, RRID AB_2744655), and Bassoon (1:800, Abcam ab82956, AB_1860018) by using antibodies. The following secondary antibodies were applied for 1 hr at RT: Alexa 488-conjugated anti-guinea pig (AB_2534117), Alexa 555-conjugated anti-mouse (AB_2535769), and Alexa-647-conjugated anti-rabbit (AB_2535813) to detect homer, bassoon, and VGLUT2, respectively; Alexa 488-conjugated anti-rabbit (AB_2762833), Alexa 555-conjugated anti-mouse IgG2a (AB_2535776), and Alexa-647-conjugated anti-mouse IgG1 (AB_2535809) to detect homer, bassoon, and VGLUT1, respectively.

Images were acquired using a Nikon CSU-W1 super resolution by optical reassignment (SoRa) spinning disk confocal microscope with a 10x objective for thalamus overview images and a 60x objective with 4x SoRa disk (27 nm pixel size) for near super-resolution synaptic images. Fluorescence signals were excited using 488, 561, and 640 nm lasers and detected using GFP, RFP, and Cy5 filters and a Hamamatsu back-thinned Orca Fusion CMOS camera. Z-stacks were acquired with Nikon Elements software with a 0.15 μm interval. Three 60x images from different fields of view (FOV) within the VPL and VPM thalamus were taken for each mouse. Images were processed with Nikon Elements Denoising software and Blind Deconvolution algorithm. All further image processing and analysis was performed using Fiji on a single z-plane from each image series.

### Image analysis

The SynapseJ plugin was used to detect glutamatergic synapses defined as overlapping homer and bassoon puncta as well as VGLUT1- and VGLUT2-positive bassoon puncta to label CT and sensory inputs, respectively (Moreno Manrique et al. 2021; Topolski et al. 2025). The minimum and maximum threshold intensities to detect each synaptic marker as separate particles were set empirically for each imaging session. The minimum and maximum puncta size parameters, set at 0.02 and 5 for both bassoon and homer, were kept constant across all images. VGLUT-positive synapses were defined as bassoon puncta overlapping VGLUT1 or VGLUT2 signal with at least one pixel at the maximum intensity threshold. VGLUT2 is highly expressed in sensory axon terminals, but also visible in VPL and VPM neurons, so to be sure we specifically detected bassoon puncta overlapping sensory terminals we performed a median blur on VGLUT2, which ensured thresholding eliminated weak VGLUT2 signal in VPL and VPM neurons and maintained strong VGLUT2 signal in axon terminals. The data from three FOV were averaged for each mouse. Experimenters were blind to genotype during imaging and analysis.

### Statistical analysis

*A priori* power analyses were performed in GPower 3.1 to estimate required sample sizes given appropriate statistical tests with α = 0.05, power (1 – β) = 0.8, and a moderate effect size or effect size based on pilot data. By using GraphPad Prism 9.4.1 statistical analysis was performed. Normality was tested by the D’Agostino-Pearson test and equal variances was tested by the Spearman’s tests with the results reported in **Supplemental Table 1**. Normal datasets with equal variances were analyzed by two-way ANOVA with Sidak’s corrections for multiple comparisons; these data were reported in the text and plotted as the group mean with standard deviation (*SD*). All frequency data for synaptic currents had log-normal distributions and most exhibited unequal variances; therefore, these data were log-transformed for statistical analysis. If any frequency data point was ≤ 1.0 Hz, then a constant value = 1.0 was added to each data point so that log transformed values were greater than zero. All transformed datasets passed normality and equal variances tests (**Supplemental Table S1**) and were analyzed by two-way ANOVA with pairwise Sidak’s tests to correct for multiple comparisons. For transformed datasets, the geometric mean of the raw (untransformed) data was reported in the text with the 95% confidence interval [95 CI], while both the raw data with geometric mean [95 CI] and log-transformed data with mean [95 CI] were plotted in the figures. These values are also reported in **Supplemental Table S2** along with group summary data including mean (SD), geometric mean [95 CI], and mean of the log-transformed data [95 CI]. ANOVAs and pairwise comparisons were only performed on the transformed data and are reported with the plots of transformed data. The ANOVA test statistics, *p* values, and degrees of freedom were reported in **Supplemental Tables S3** and **S4**, and pairwise Sidak’s test *p* values were reported in the figures and/or figure legends.

## RESULTS

### DS mice have reduced glutamatergic input to VPL neurons after seizure onset and in adulthood

To investigate glutamatergic synapse dysfunction in VPL and VPM neurons, we first measured the amplitude and frequency of sEPSCs in acute brain slices from mice aged approximately 2 weeks (2W, P13-P17, before highest seizure burden), 4 weeks (4W, P28-P32, after seizure onset), and 8 weeks (8W, P58-P62, adult; **Figure 1A-C**). The mean frequency of VPL sEPSCs in DS mice was similar to WT in 2W mice (DS: 3.2 Hz [2.1, 5.0]; WT: 3.0 Hz [2.3, 4.0]) and 8W mice (DS:2.5 Hz [1.6, 3.9]; WT: 3.8 Hz [2.7, 5.4), but was significantly reduced compared to WT in 4W mice (DS: 2.1 Hz [1.6, 2.9]; WT: 3.6 Hz [2.7, 4.9]) **Figure 1D-F).** Whereas, mean sEPSC amplitude was similar between WT and DS mice for VPL neurons at all ages (2W: DS 16.2 pA, *SD* 0.9; WT 16.9 pA, *SD* 2.6; 4W: DS 15.0 pA, *SD* 3.5; WT: 17.7 pA, *SD* 4.3; 8W: DS 17.4 pA, *SD* 3.0; WT 15.9 pA, *SD* 2.6; **Figure 1G**). In VPM neurons, mean sEPSC frequency was similar between DS and WT mice at 2W (WT: 1.6 [1.2, 1.9]; DS: 1.6 Hz [1.1, 2.2]), 4W (DS: 2.2 Hz [1.7, 2.9]; WT: 2.3 Hz [1.6, 3.4]), and 8W (DS: 2.3 Hz [1.5, 3.6]; WT: 2.1 Hz [1.5, 3.0]; **Figure 1H-J**). Likewise, VPM sEPSC amplitude in DS mice did not differ from WT in all three age groups (2W: DS 14.0 pA, *SD* 2.3; WT 14.2 pA, *SD* 1.5; 4W: DS 16.0 pA, *SD* 2.9; WT: 17.4 pA, *SD* 4.2; 8W: DS 17.3 pA, *SD* 2.6; WT 16.7 pA, *SD* 2.7; **Figure 1K**). These data suggest that reduced spontaneous glutamatergic input to VPL neurons in DS mice appears after seizure onset and normalizes by adulthood, while spontaneous glutamatergic input to VPM neurons is unaffected in DS mice.

**Figure 1.**
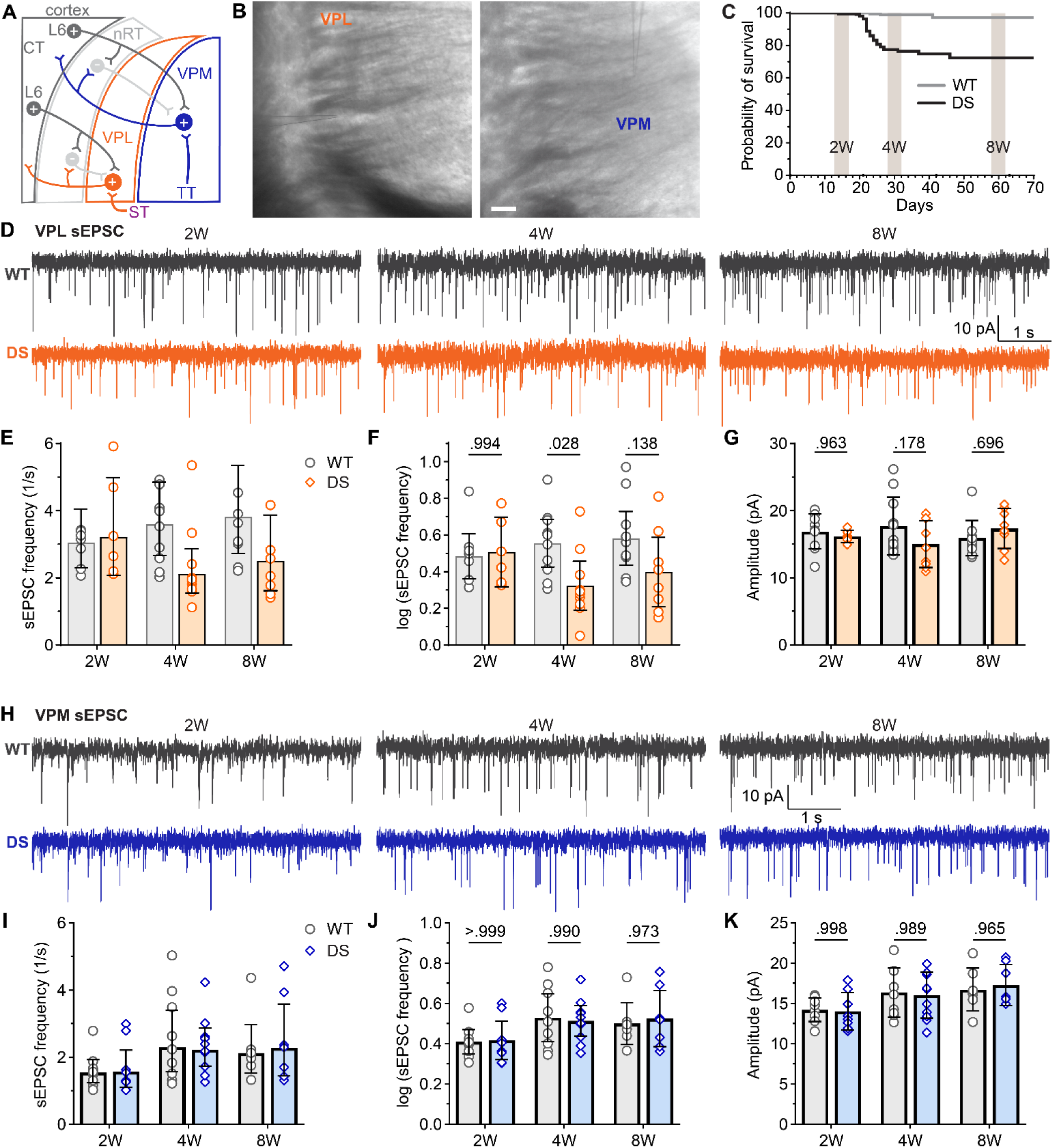
VPL neuron sEPSC frequency is reduced in DS mice after seizure onset. **A**. Circuit diagram shows excitatory and inhibitory synaptic inputs to VPL and VPM neurons (ST: spinothalamic tract; TT: trigeminothalamic tract). **B.** Images of VPL and VPM recordings in acute thalamic slices**. C.** A survival timeline for WT and DS mice illustrates the timing of three age groups in this study: 2W, 4W, and 8W relative to the period of highest seizure burden (P17-P24). **D.** Representative voltage-clamp recordings from VPL neurons show sEPSCs in WT and DS mice aged 2W (WT: n = 9; DS: n = 6), 4W (WT: n = 10; DS: n = 9), and 8W (WT: n = 11; DS: n = 8). **E.** The mean sEPSC frequency data were plotted as raw values with the geometric mean ± 95CI, which had a non-normal distribution, and **F**) transformed log_10_ values with group mean ± 95CI, which were used for statistical analyses. **G.** Mean sEPSC amplitudes were plotted for each mouse with the group mean ± *SD*. **H.** Representative VPM recordings show sEPSCs in WT and DS mice aged 2W (WT: n = 9; DS: n = 8), 4W (WT: n = 9; DS: n = 11), 8W (WT: n = 7; DS: n = 7). **I-K**. The mean sEPSC frequency and amplitude data were plotted as described for VPL. The data were analyzed by two-way ANOVA with pairwise Sidak’s comparisons (p-values shown in plots) between WT and DS at each age.

The observed differences in sEPSC frequency could have been due to varying activity levels in presynaptic axons, probability of release, and/or synapse number. To further understand how synaptic transmission was affected, we measured mEPSC frequency and amplitude in the absence of action potential driven synaptic release (**Figure 2A,E**). In 2W mice, we found no significant changes in mean mEPSC frequency in VPL (WT: 3.3 Hz [2.3, 4.8]; DS: 3.3 Hz [2.5, 4.3]; **Figure 2B,C**) or VPM neurons (WT: 1.6 Hz [.12, 2.2]; DS: 1.9 Hz [1.3, 2.8]; **Figure 2F,G**). However, in 4W mice, the mean frequency of VPL mEPSCs was reduced in DS mice (1.8 Hz [1.5, 2.3]) compared to WT (3.5 Hz [2.8, 4.4]), with no change in mean VPM MEPSC frequency (DS: 1.8 Hz [1.4, 2.4]; WT: 1.8 Hz [1.3, 2.7]). In adult mice, VPL neuron mEPSC frequency remained significantly reduced in DS (2.0 Hz [1.7, 2.6]) relative to WT (3.2 Hz [2.1, 4.9]), while VPM mEPSC frequency was similar between DS (2.4 Hz [1.7, 3.5]) and WT (2.1 Hz [1.5, 3.1]) mice. The mean mEPSC amplitude was reduced in DS mice relative to WT at 4W (DS 14.3 pA, *SD* 1.9; WT 17.7 pA, *SD* 1.6), but not at 2W (DS 13.1, *SD* 1.9; WT 12.1 pA, *SD* 1.8) or 8W (DS 18.6 pA, *SD* 2.9; WT 17.1 pA, *SD* 2.8; **Figure 2D**). Mean mEPSC amplitude did not differ between DS and WT for VPM neurons in all age groups (2W: DS 14 pA, *SD* 1.8; WT: 12.5 pA, *SD* 2.3; 4W: DS 13.1 pA, *SD* 2.0; WT: 13.9 pA, *SD* 2.6; 8W: DS 12.9 pA, *SD* 2.0; WT 13.9 pA, *SD* 2.9; **Figure 2H; Supplementary Table 1)**. Altogether, these data suggest that specifically VPL neurons, but not VPM neurons, have a reduced number of glutamatergic inputs or reduced probability of glutamate release, which appears after seizure onset in DS mice and is maintained into adulthood.

**Figure 2.**
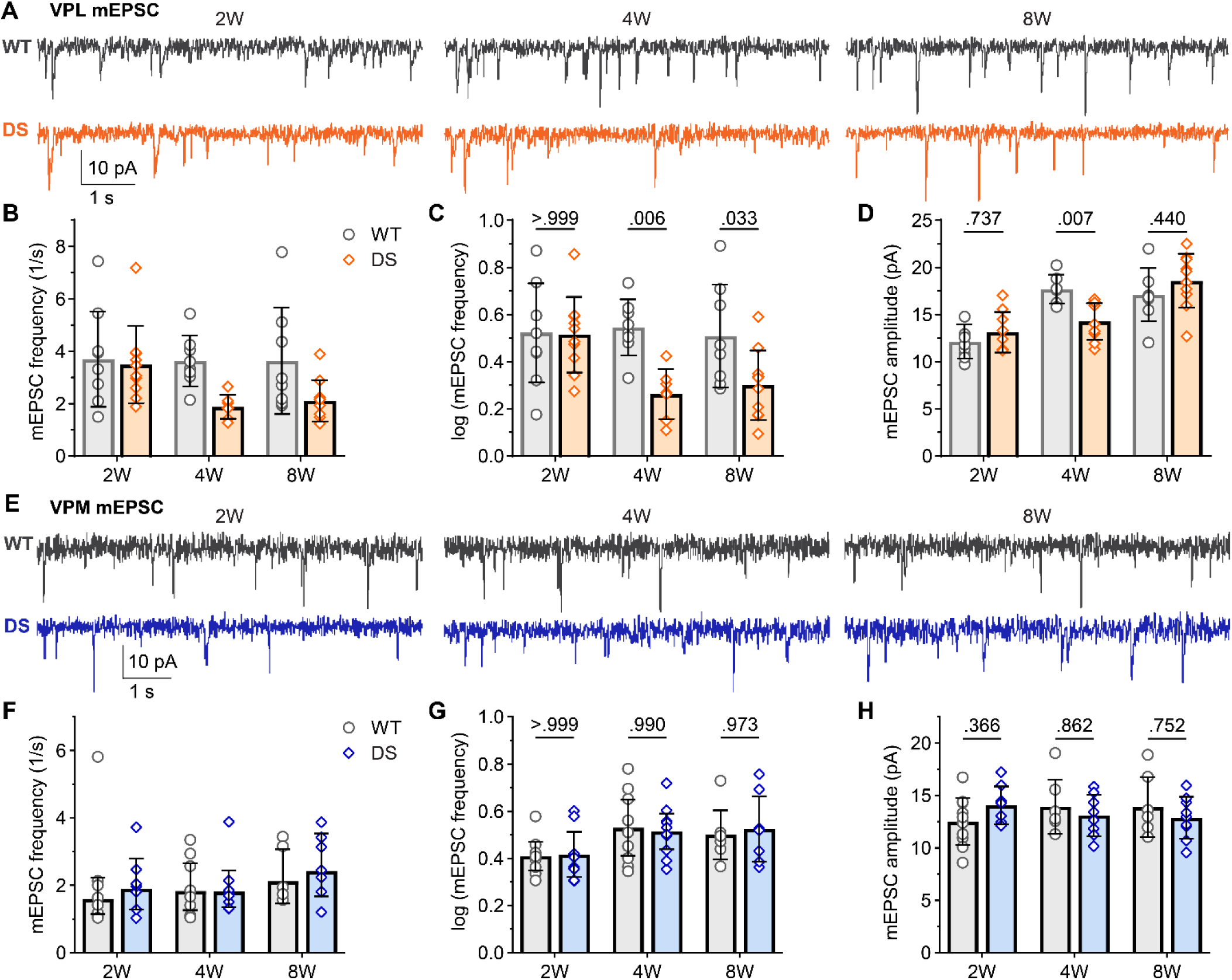
DS mice show reduced mEPSC frequency in VPL neurons after seizure onset. **A.** Representative voltage-clamp recordings from VPL neurons show mEPSCs in WT and DS mice aged 2W (WT: n = 9; DS: n = 10), 4W (WT: n = 8 ; DS: n = 7), 8W (WT: n = 8; DS: n = 10). The mean mEPSC frequency was plotted as raw data (geometric mean ± 95CI) and **C**) transformed log_10_ values (mean ± 95CI) as the data had a non-normal distribution. **D.** Mean mEPSC amplitude were plotted for each mouse with the group mean ± *SD*. **E.** Representative VPM recordings show mEPSCs in WT and DS mice aged 4W (WT: n = 11; DS: n = 7), 4W (WT: n = 8; DS: n = 8), 8W (WT: n = 6; DS: n = 7). **F-H**. The mean mEPSC frequency and amplitude data were plotted as described for VPL. VPL and VPM data were analyzed by two-way ANOVA with pairwise Sidak’s comparisons (p-values shown on plots) between WT and DS at each age.

### Sensory synapse number is selectively reduced in VPL neurons in DS mice

VPL and VPM neurons receive descending glutamatergic input from CT neurons and ascending glutamatergic input carrying sensory information from the spinal cord and brainstem, respectively. To determine whether one or both of these inputs were affected in DS mice, we quantified the number of CT and sensory synapses in the VPL and VPM utilizing immunohistochemistry and high-resolution imaging. Bassoon immunostaining was used to identify presynaptic release sites, and CT inputs were labeled by immunostaining for VGLUT1 (**Figure 3A,B**), while sensory terminals were labeled with VGLUT2 immunostaining (**Figure 4A,B**). We compared the number of VGLUT1- and VGLUT2-positive bassoon puncta to approximate CT and sensory synapse density in WT and DS mice. The number of VGLUT1-positive bassoon puncta in the VPL was not significantly different between WT and DS mice at 2W (WT: 907, *SD* 217; DS: 1042, *SD* 197), 4W (WT: 1772, *SD* 311; DS: 1631, *SD* 312), or 8W (WT: 1637, *SD* 170; DS: 1508, *SD* 281; **Figure 3C,D; Supplemental Figures S1 and S2**). We also detected similar numbers of VGLUT1-positive bassoon in VPM neurons across WT and DS mice at 2W (WT: 785, *SD* 176; DS: 1069, *SD* 145), 4W (WT: 1346, *SD* 286; DS: 1327, *SD* 283), or 8W (WT: 1335, *SD* 187; DS: 1479, *SD* 119; **Figure 3C,E**). The numbers of VGLUT2-positive bassoon puncta in the VPL neurons were similar between WT and DS mice aged 2W (WT: 148, *SD* 30; DS: 135, *SD* 30), but were reduced in 4W DS mice (WT: 187, *SD* 34; DS: 123, *SD* 25) and 8W DS mice (WT: 191, *SD* 46; DS: 117, *SD* 35) compared to WT (**Figure 4C,D; Supplemental Figures S3 and S4)**. No significant differences in the number of VGLUT2-positive bassoon puncta were detected in the VPM between WT and DS mice across all three age groups (2W: WT 97, *SD* 30; DS 113, *SD* 27; 4W: WT 108, *SD* 16; DS 101, *SD* 17; 8W: WT 107, *SD* 29; DS 112, *SD* 27; **Figure 4C,E).**

**Figure 3.**
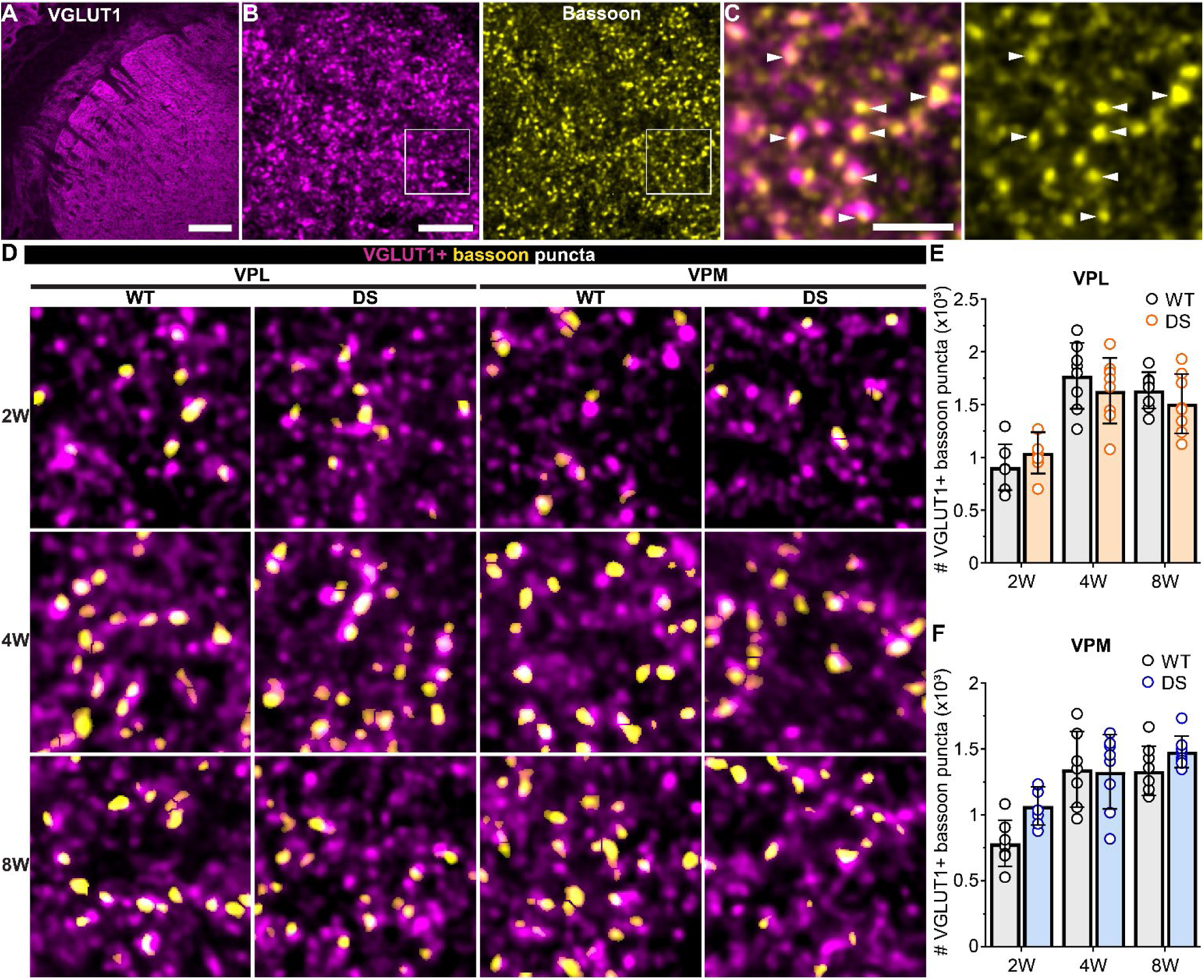
VGLUT1-positive CT synapse numbers are unaffected in VPL and VPM thalamus of DS mice. **A.** 10x image of the thalamus including the VPL and VPM immunostained for VGLUT1. **B**. 60x images maximum intensity projection of VGLUT1 and bassoon staining in the VPM thalamus. **C.** Boxed region from panel B shows individual bassoon puncta (arrowheads) overlapping VGLUT1 signal in a single z-plane. Scale: A) 200 µm, B) 5 µm, and C) 2 µm. **D.** Representative ROIs from 60x VPL and VPM images show VGLUT1-positive bassoon puncta in WT and DS mice aged 2W, 4W, and 8W. The number of VGLUT1-positive bassoon puncta were averaged across three images from the **E)** VPL [2W (WT: n = 7; DS: n = 7), 4W (WT: n = 8 ; DS: n = 8), and 8W (WT: n = 8; DS: n = 8 )] and **F)** VPM [2W (WT: n = 7; DS: n = 7), 4W (WT: n = 7; DS: n = 8 ), and 8W (WT: n = 8 ; DS: n = 8 ;)] in each mouse. The group mean and SD were plotted with data points from all mice. Data were analyzed by one-way ANOVA with posthoc Sidak’s tests between genotypes at each age (p-values shown in plots).

**Figure 4.**
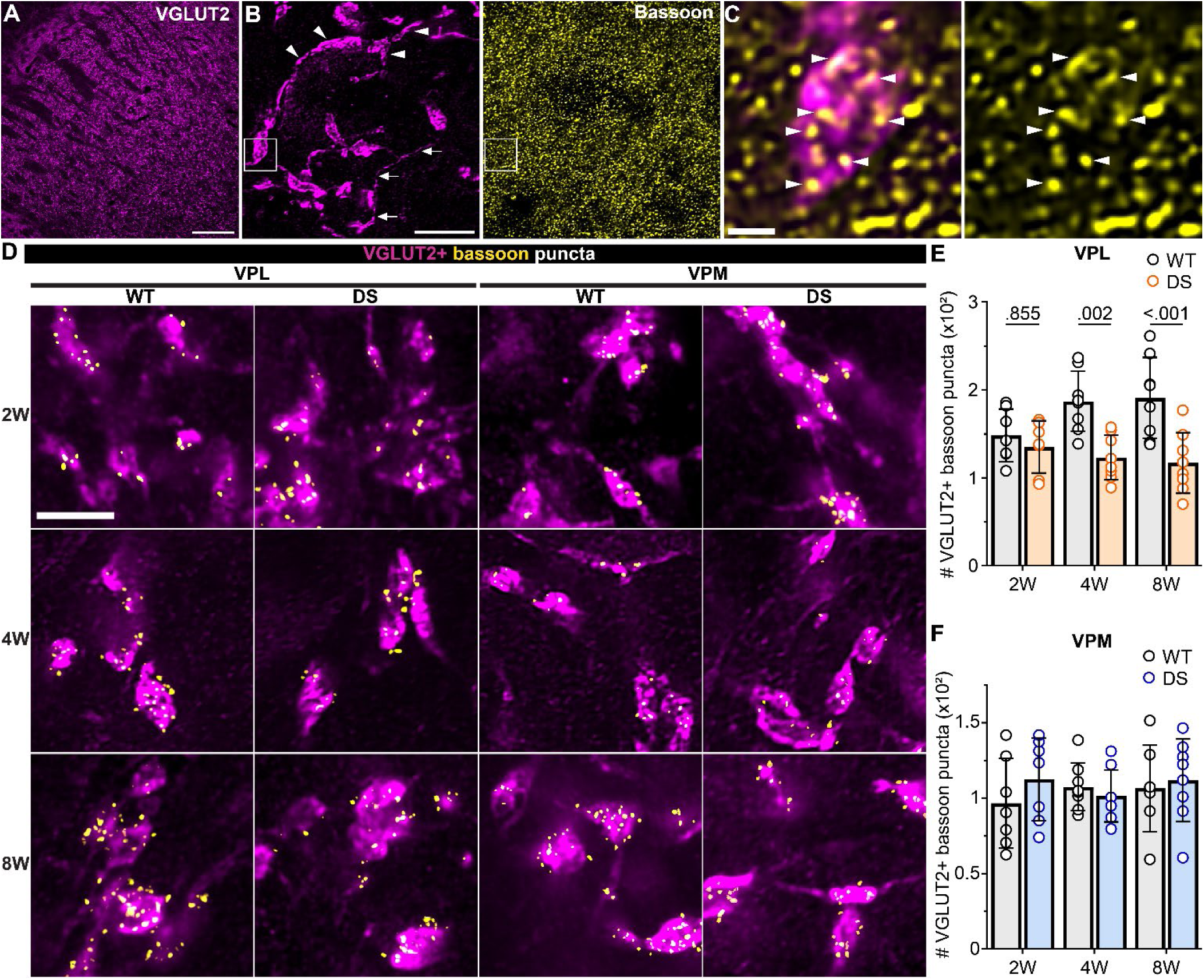
VGLUT2-positive synapse number in the VPL thalamus is reduced in DS mice. **A.** 10x image of the thalamus including the VPL and VPM immunostained for VGLUT2. **B**. 60x images maximum intensity projection of VGLUT2 and bassoon staining in the VPM thalamus. Arrows highlight a single sensory axon with a large VGLUT2-positive terminal. Arrowheads highlight VGLUT2-postive presynaptic terminals on the soma and prozimal dendrites of a VPM thalamus neuron. C. Boxed region from panel B shows individual bassoon puncta (arrowheads) overlapping VGLUT2 signal in a sensory axon terminal in a single z-plane. Scale: A) 200 µm, B) 10 µm, and C) 1 µm . D. Representative regions of interest (ROIs) from 60x VPL and VPM images show VGLUT2-positive bassoon puncta in WT and DS mice aged 2W, 4W, and 8W. The number of VGLUT2-positive bassoon puncta were averaged across three images from the **E)** VPL [2W (WT: n = 7; DS: n = 7 ), 4W (WT: n = 8 ; DS: n = 8), and 8W (WT: n = 8 ; DS: n =8)] and **F)** VPM [2W (WT: n = 7 ; DS: n = 7), 4W (WT: n = 8 ; DS: n = 8), and 8W (WT: n = 7 ; DS: n = 8)] in each mouse. The group mean and SD were plotted with data points from all mice. Data were analyzed by one-way ANOVA with posthoc Sidak’s tests between genotypes at each age (p-values shown in plots).

We also quantified total glutamatergic synapses in VPL and VPM, with single synapses identified as overlap-ping bassoon and homer (postsynaptic) puncta. The total number of synapses was not significantly different across WT and DS mice in any age group as measured by quantifying all overlapping pairs of homer and bassoon in VPL neurons (2W: WT 943, *SD* 189; DS 908, *SD* 133; 4W: WT 1706, *SD* 216; DS 1498, *SD* 277; 8W: WT 1637, *SD* 275; DS 1568, *SD* 280) or VPM neurons (2W: WT 995, *SD* 172; DS 923, *SD* 154; 4W: WT 1535, *SD* 199; DS 1417, *SD* 210; 8W: WT 1396, *SD* 258; DS 1267, *SD* 201; **Supplemental Figures S5 and S6**). These data are consistent with our CT and sensory synapse quantification, because sensory synapses make up a small portion (<10%) of total glutamatergic synapses in somatosensory thalamus; thus, any change in this population would not necessarily be detectable when assessing the whole population of glutamatergic synapses. Although few in number, sensory inputs are large, proximal synapses with multiple release sites with high release probability that contribute a substantial portion of the total excitatory input to VPL and VPM neurons. Altogether, these data suggest that show a cell-type- and input-specific reduction of sensory synapses onto VPL neurons.

As a second means of assessing whether one or both of these inputs were affected, we took advantage of the known input-specific differences in EPSC decay time in thalamocortical neurons with sensory EPSCs having a substantially faster decay time than cortical EPSCs (Studtmann et al., 2022). We plotted decay times for all mEPSCs from each neuron as a frequency histogram, and separated the resulting bimodal distribution at a mathematically determined critical value **(Supplemental Figure S7)**. Then we analyzed the amplitude and frequency of putative sensory (Type 1) and cortical (Type 2) mEPSCs to determine if differences observed in the total mEPSC population resulted from one or both input types. In VPL neurons, the frequency of Type 1 mEPSCs at 2W was not different between WT (1.2 Hz [0.8, 1.6]) and DS (1.0 [0.8, 1.3]) mice (**Figure 5A,B**). Notably, the Type 1 mEPSC frequency was significantly reduced in DS mice at 4W (0.8 Hz [0.6, 0.9]) compared to WT (1.9 Hz [1.5, 2.6]), and this reduction was maintained in adult mice aged 8W (WT: 1.6 Hz [0.9, 2.9]; DS: 1.0 [0.8, 1.4]). Interestingly, Type 2 mEPSC frequency was not statistically different between WT and DS mice at 2W (WT: 1.8 [1.2, 2.7]; DS: 1.8 Hz [1.3, 2.4]), 4W (WT: 1.7 Hz [1.3, 2.2]; DS: 1.2 Hz [0.9, 1.7]), or 8W (WT: 1.5 Hz [1.0, 2.5]; DS: 1.4 Hz [1.2, 1.7]). The amplitude of Type 1 mEPSCs in VPL neurons were significantly reduced in 4W DS mice (13.5 pA, *SD* 1.5) relative to WT (16.1 pA, *SD* 2.5), but not at 2W (WT: 12.4 pA, *SD* 1.3; DS: 11.7 pA, *SD* 1.4) or 8W (WT: 14.8 pA, *SD* 2.1; DS: 16.4 pA, *SD* 2.1; **Figure 5C).** There were no statistically significant differences in Type 2 mEPSC amplitude in VPL neurons at 2W (WT: 14.2 pA, *SD* 2.5; DS: 12.9 pA, *SD* 2.1), or 8W (WT: 16.9 pA, *SD* 2.9; DS: 18.7 pA, *SD* 2.5). Altogether, these electrophysiology and synapse imaging data suggest that VPL neurons display reduced glutamatergic transmission in a synapse-specific manner leading due to a putative reduction in ascending sensory input, while cortical input remains unchanged. There were no changes in the amplitude or frequency of Type 1 and Type 2 mEPSCs in VPM neurons further suggesting that this reduced glutamatergic input to somatosensory thalamus is cell-type-specific (**Supplemental Figure S8; Supplemental Table S2)**.

**Figure 5.**
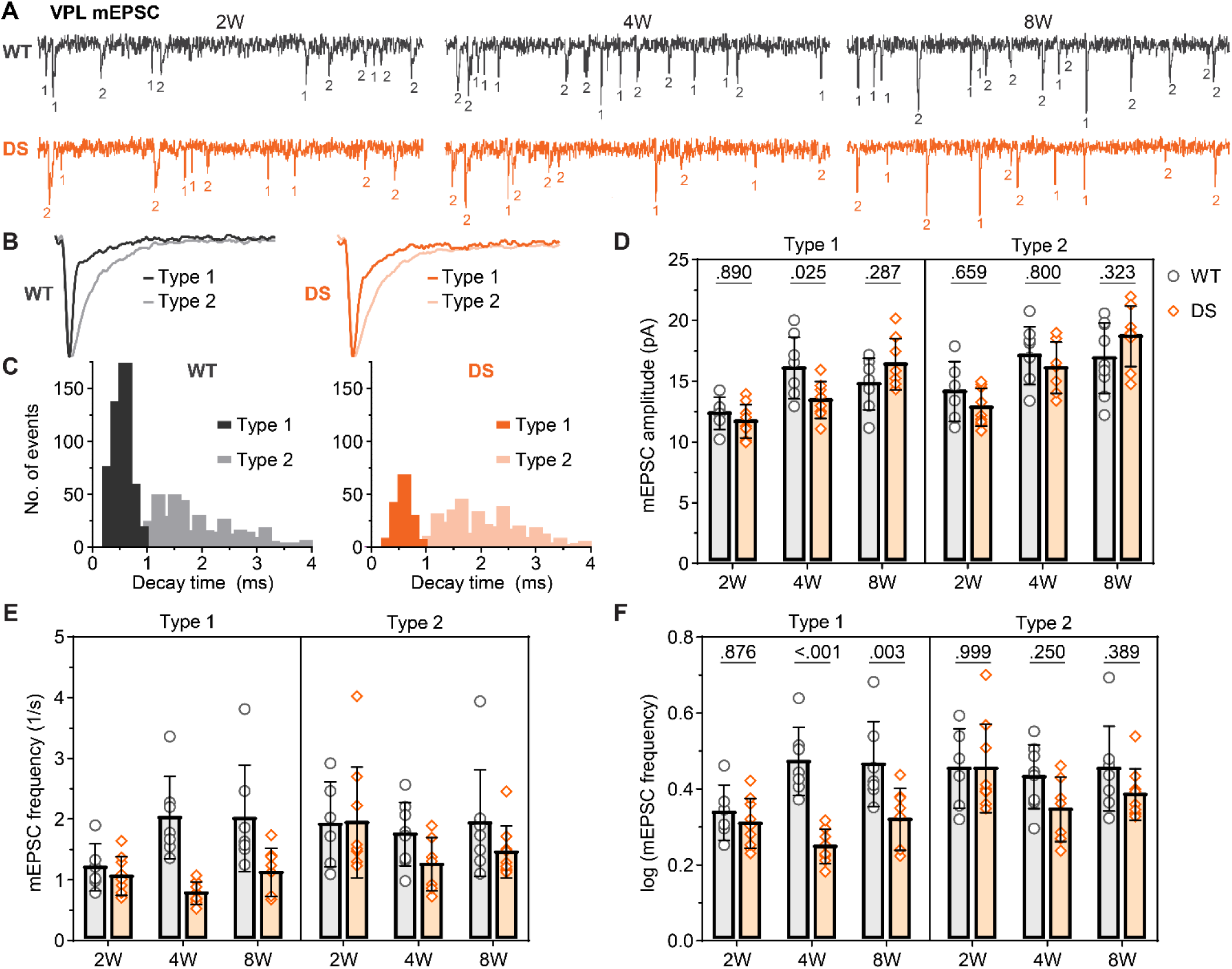
DS mice show a selective reduction in Type 1 mEPSC frequency in VPL neurons after seizure onset. **A.** Representative voltage-clamp recordings from VPL neurons show mEPSCs in WT and DS mice aged 2W (WT: n = 6; DS: n = 9), 4W (WT: n = 7; DS: n = 8), and 8W (WT: n = 7; DS: n = 8) with each mEPSC denoted as Type 1 (fast, putative sensory) or Type 2 (slow, putative CT) based on their decay kinetics. B. Representative ensemble average traces show Type 1 and Type 2 mEPSCs from WT and DS mice. **C.** Histograms show representative bimodal frequency distributions of mEPSC decay kinetics for all mEPSCs from representative WT and DS recordings. **D.** The mean mEPSC frequency was plotted as raw data (geometric mean ± 95CI) and **E**) transformed log10 values (mean ± 95CI) as the data had a non-normal distribution. **D.** Mean mEPSC amplitude were plotted for each mouse with the group mean ± *SD*. The data were analyzed by two-way ANOVA with pairwise Sidak’s comparisons (p-values shown on plots) between WT and DS at each age.

### DS mice show reduced inhibitory synaptic transmission in VPL neurons, but not VPM neurons

VPL and VPM neurons receive dense inhibitory input from nRT neurons, which show reduced tonic firing and excess burst firing in DS mice (Hedrich et al., 2014; Ritter-Makinson et al., 2019; Studtmann et al., 2022). To determine how inhibitory synaptic transmission is affected across development in DS, we measured spontaneous inhibitory postsynaptic currents (sIPSCs) across the three age groups (**Figure 6**). At 2W, we found no significant changes in mean sIPSC frequency in VPL (WT: 6.1 Hz [4.3, 8.8]; DS: 7.3 Hz [5.4, 10.0]); however, DS mice showed reduced sIPSC frequency at compared to WT mice at 4W (WT: 7.3 Hz [5.2, 10.1]; DS: 3.7 Hz [2.6, 5.2]) and 8W (WT: 8.2 Hz [6.0, 11.4]; DS: 4.8 Hz [3.6, 6.4]; **Figure 6A-C**)). Mean sIPSC amplitude was similar for VPL neurons at 2W (WT: 25.3 pA, *SD* 4.3; DS: 25.2 pA, *SD* 5.7), 4W (WT: 25.8 pA, *SD* 5.4, DS: 26.8 pA, *SD* 6.2), and 8W (WT: 29.9 pA, *SD* 4.3, DS: 27.3 pA, *SD* 3.0 ; **Figure 6D**). These data reproduced the observed effects in 4W mice from our previous report, (Studtmann et al. 2022), and demonstrate that sIPSC frequency remains reduced in adult mice. In VPM neurons, sIPSC frequency in 4W DS mice (4.1 Hz, [3.1, 5.3]) was significantly reduced compared to WT (9.2 Hz [6.3, 13.3]; **Figure 6E-G**). VPM sIPSC frequency was similar between WT and DS mice at 2W (WT: 8.7 Hz [6.9, 11.0]; DS: 6.6 Hz [4.2, 10.3]) and 8W (WT: 7.5 Hz [4.9, 11.4]; DS: 6.6 Hz [4.7, 9.2]). Mean sIPSC amplitude in VPM neurons did not differ between WT and DS mice at 2W (WT: 27.1 pA, *SD* 6.6; DS, 22.1 pA, *SD* 2.6), 4W (WT: 28.5 pA, *SD* 6.2; DS: 26.1 pA, *SD* 4.6), or 8W (WT: 28.8 pA, *SD* 1.6; DS: 28.7 pA, *SD* 7.7; **Figure 6H**). These data suggest that DS mice have reduced spontaneous inhibitory synaptic input that arises after seizure onset in both VPL and VPM neurons, but is only maintained into adulthood in VPL neurons.

**Figure 6.**
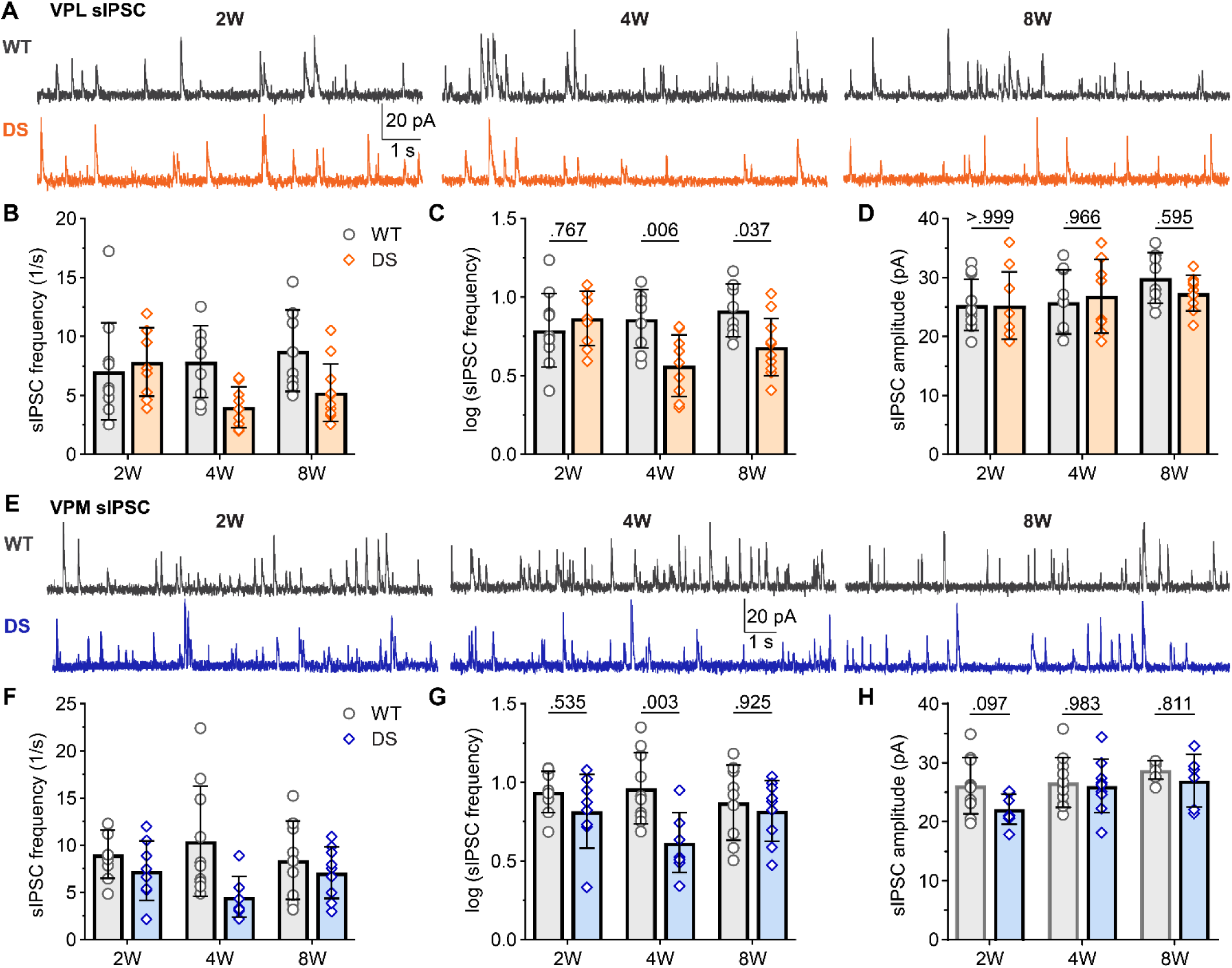
Reduced sIPSC frequency persists in adult DS mice in VPL neurons, but normalizes in VPM neurons. **A.** Representative voltage-clamp recordings from VPL neurons show sIPSCs in WT and DS mice aged 2W (WT: n = 11; DS: n = 9), 4W (WT: n = 9; DS: n = 8), and 8W (WT: n = 8; DS: n = 11). **B.** The mean sIPSC frequency data were plotted as raw values with the geometric mean ± 95CI, which had a non-normal distribution, and **F**) transformed log_10_ values with group mean ± 95CI, which were used for statistical analyses. **G.** Mean sIPSC amplitudes were plotted for each mouse with the group mean ± *SD*. **H.** Representative VPM recordings show sIPSCs in WT and DS mice aged 2W (WT: n = 9; DS: n = 8), 4W (WT: n = 10; DS: n = 8), and 8W (WT: n = 7; DS: n = 6). **I-K**. The mean sIPSC frequency and amplitude data were plotted as described for VPL. The data were analyzed by two-way ANOVA with pairwise Sidak’s comparisons (p-values shown in plots) between WT and DS at each age.

We further recorded mIPSCs to investigate whether WT and DS mice exhibit any differences in synapse number or release probability that could have been obscured by differences in presynaptic neuron firing in sIPSC recordings. (**Figure 7A,E**). The data from 4W mice reproduced the observed effects from our previous report, including decreased frequency of VPL mIPSCs in DS mice (4.0 Hz [3.1, 5.3]) compared to WT (7.8 Hz [6.0, 10.2]; **Figure 7B,C**), with no change in mean VPM mIPSC frequency (DS: 6.2 Hz [3.8, 10.3]; WT: 4.9 Hz [3.6, 6.5]; **Figure 7F,G**). The mean mIPSC frequency did not differ between DS and WT at 2W for either VPL (DS: 5.5 Hz [4.0, 7.7; WT: 6.1 Hz, [4.1, 9.0]) or VPM neurons (DS: 6.3 Hz, [4.2, 9.3]; WT: 4.7 Hz [3.4, 6.5]). Similarly, there were no significant changes in mean mIPSC frequency at 8W in VPL (DS: 3.4 Hz [2.4, 4.8]; WT: 5.3 Hz [3.9, 7.4]) or VPM neurons (DS: 4.8 Hz [3.4, 6.9]; WT: 3.7 Hz [3.1, 4.4]). Mean mIPSC amplitude was similar for VPL neurons at 2W (WT: 23.2 pA, *SD* 3.6; DS: 22.0, *SD* 2.1), 4W (WT: 26.9 pA, *SD* 3.9; DS: 25.3 pA, *SD* 5.4), and 8W (WT: 25.6 pA, *SD* 3.9; DS: 25.3, *SD* 3.2).

**Figure 7.**
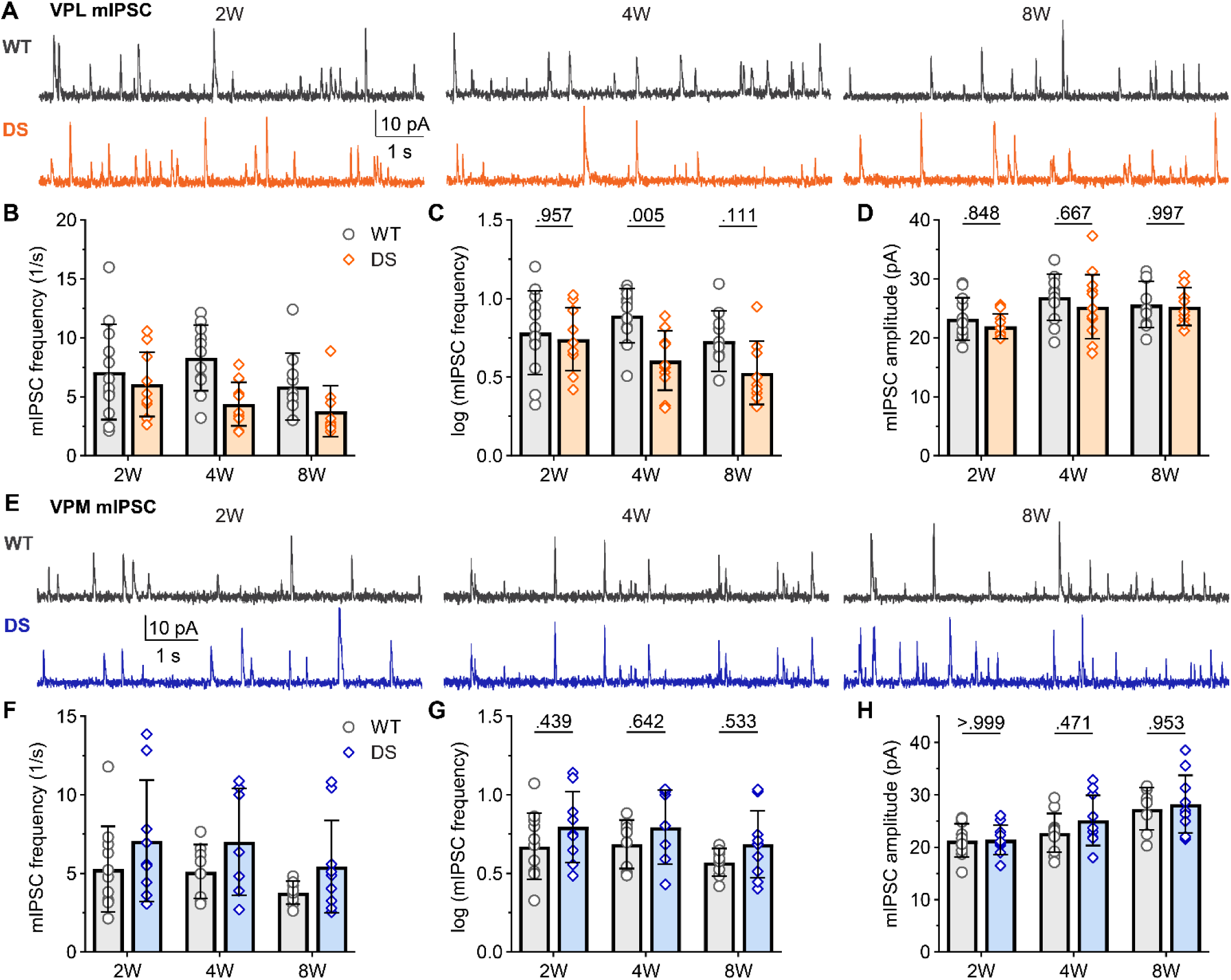
DS mice show reduced mIPSC frequency in VPL neurons after seizure onset. **A.** Representative voltage-clamp recordings from VPL neurons show mIPSCs in WT and DS mice aged 2W (WT: n = 12; DS: n = 10), 4W (WT: n = 11; DS: n = 12), and 8W (WT: n = 10; DS: n = 9). The mean mIPSC frequency was plotted as raw data (geometric mean ± 95CI) and **C**) transformed log10 values (mean ± 95CI) as the data had a non-normal distribution. **D.** Mean mIPSC amplitude were plotted for each mouse with the group mean ± *SD*. **E.** Representative VPM recordings show mIPSCs in WT and DS mice aged 2W (WT: n = 11; DS: n = 9), 4W (WT: n = 9; DS: n = 7), and 8W (WT: n = 8; DS: n = 10). **F-H**. The mean mIPSC frequency and amplitude data were plotted as described for VPL. VPL and VPM data were analyzed by two-way ANOVA with pairwise Sidak’s comparisons (p-values shown on plots) between WT and DS at each age.

Likewise, mIPSC amplitude was similar between WT and DS mice in VPM neurons at 2W (WT: 21.3 pA, *SD* 3.2; DS: 21.5 pA, *SD* 2.8), 4W (WT: 22.8 pA, *SD* 3.7; DS: 25.1, *SD* 4.8), and 8W (WT: 27.3 pA, *SD* 4.1; DS: 28.2, *SD* 5.5).

Together, these data show that both cell types exhibit reduced frequency of spontaneous inhibitory input, but this deficit is only maintained through adulthood in the VPL thalamus. Whereas, the strength of inhibitory synapses does not differ between WT and DS mice in either cell type across the age groups.

## DISCUSSION

This study reveals a selective and persistent reduction of both excitatory and inhibitory input to VPL thalamus in a mouse model of DS. In contrast, excitatory input to VPM neurons remains largely unaffected and inhibitory input is transiently impaired in DS mice, underscoring a region-specific vulnerability within the somatosensory thalamus. Our findings further indicate that these impairments emerge after seizure onset, providing new insight into the time course of altered synaptic transmission and thalamus dysfunction in DS.

We uncovered a significant reduction in both spontaneous and miniature EPSC frequency in VPL neurons in DS mice at 4 and 8 weeks of age, with amplitude reduced only in 4 week-old mice. These data suggest a decrease in the number of glutamatergic synapses and/or a reduced probability of glutamate release. Immunohistochemical analysis confirmed a reduction specifically in VGLUT2-positive sensory synapses in VPL neurons at 4 and 8 weeks of age, while VGLUT1-positive CT synapses remain unchanged. This finding was further supported by reduced frequency of mEPSC frequency at putative sensory inputs (Type 1, faster decay time), while putative CT inputs (Type 2, slower decay time) were not significantly affected. Notably, synapse number and function appeared normal in 2-week-old DS mice, indicating that there is selective vulnerability of sensory inputs to VPL neurons that emerges after the period of highest seizure burden at age P17 – P24.

Na_V_1.1 expression increases rapidly during the second week of life in mice (Cheah et al. 2013; Gordon et al. 1987; Scheinman et al. 1989; Beckh et al. 1989), and thus DS mice with Na_V_1.1 haploinsufficiency develop seizures beginning in the third week (Yu et al. 2006b; Miller et al. 2014). Na_V_1.1 haploinsufficiency could impact neuronal excitability and subsequently disrupt synapse development before seizure onset; however, based on our results, it is likely that altered thalamic synapse development is a consequence of circuit hyperexcitability and/or seizures. Synapse number was reduced in adult DS mice, whereas synapse strength normalized by this age, indicating that altered synapse maintenance may affect excitatory synaptic transmission more than altered function of individual synapses in adult DS mice. However, we cannot rule out the existence of changes in release probability at somatosensory synapses to VPL neurons. These findings align with a previous study showing deficits to sensory inputs to VPL neurons in 4-week-old DS mice (Studtmann et al. 2022) and add to this framework by demonstrating that sensory input deficits develop after seizure onset and are maintained into adulthood.

Importantly, inhibitory input to VPL neurons was also impaired in DS mice, developing after seizure onset and persisting until adulthood. The combined loss of excitatory sensory input and inhibitory control in VPL neurons suggests a profound disruption of thalamocortical processing in DS. VPL neurons integrate ascending sensory signals from the body and send projections to primary somatosensory cortex, playing a key role in tactile perception and sensorimotor integration. The loss of VGLUT2-positive sensory synapses, which are large and proximal with multiple release sites, likely results in diminished excitatory drive. Concurrently, reduced inhibitory input from the nRT may impair the temporal precision and gain control of thalamic output (Ritter-Makinson et al. 2019; Iavarone et al. 2023). This dual synaptic impairment persists through adulthood and may contribute to the sensory abnormalities observed in DS patients, including altered pain perception and sensorimotor integration deficits (Guzzetta 2011; Anwar et al. 2019; Turner et al. 2017; Domaradzki and Walkowiak 2023). This finding also supports the hypothesis that synaptic dysfunction in the thalamus could be a central feature of DS pathophysiology, extending beyond seizures to encompass broader cognitive and behavioral impairments that persist in adults after seizures have largely waned. However, a limitation of this study is that it remains unknown how these synaptic alterations impact thalamocortical output from the VPL in response to sensory excitatory drive, which will be critical for understanding how sensory information flow through the thalamus is affected in DS.

Interestingly, VPM synaptic input was largely unaffected in DS mice at any age. The cell-type-specificity of synaptic dysfunction in the somatosensory thalamus may reflect differences in the developmental trajectory, connectivity, or intrinsic properties of VPL versus VPM neurons. Indeed, VPL neurons have been shown to exhibit stronger excitatory input and distinct firing properties compared to VPM neurons (Studtmann et al. 2023) and receive somewhat distinct inhibitory input from the nRT (Clemente-Perez et al. 2017). Moreover, Na_V_1.1 haploin-sufficiency impacts the two cell types differently, with VPL neurons having reduced excitability and VPM neurons show a modest increase in excitability (Studtmann et al. 2022). Thus, it is possible that the distinct activity levels in these two cell populations contribute to different synaptic phenotypes. Alternatively, distinct connectivity could contribute to cell-type-specific synaptic alterations in DS mice as presynaptic sensory neurons innervating VPL and VPM could be affected differently. Both dorsal root ganglia (DRG) and trigeminal ganglia sensory neurons express Na_V_1.1 (Fukuoka et al. 2008; Ho and O’Leary 2011), and Na_V_1.1 expression in these cells is important for mechanosensation and proprioception (Osteen et al. 2016; Salvatierra et al. 2018; Pineda-Farias et al. 2021). However, the functional consequences of Na_V_1.1 haploinsufficiency have not been directly compared between DRG and trigeminal sensory neurons. Additionally, VPL and VPM neurons receive input from distinct nRT neuron populations that could be impacted differently in DS, further contributing to cell-type-specific thalamic dysregulation in DS (Clemente-Perez et al. 2017). Altered sensory processing and sleep deficits are key behavioral phenotypes in patients with loss-of-function Na_V_1.1 patients throughout their lifespan (Ding et al. 2021; Lossin 2009). Determining which circuit components are directly impacted by Na_V_1.1 haploinsufficiency versus those altered by circuit hyperexcitability will be required for a better understanding of how ascending sensory input versus central sensory information processing are impacted across disease stages.

These findings underscore the importance of determining how impaired excitatory and inhibitory synaptic integrity in the VPL contribute to thalamocortical dysfunction and whether correcting these impairments is beneficial for behavioral symptoms in DS. Further studies are needed to elucidate the molecular mechanisms underlying vulnerability of sensory synapses relative to CT synapses. It will be important to determine whether these differences establish vulnerability only during development or continually contribute to sensory synapse dysfunction through adulthood. Therapeutic strategies aimed at preserving or restoring sensory synapse function in the VPL may hold promise for mitigating persistent DS-associated sensory symptoms in adult patients. Moreover, the differential vulnerability of VPL versus VPM neurons raises intriguing questions about the mechanisms governing synaptic resilience in distinct thalamic nuclei. Investigating the molecular and developmental factors that confer protection to VPM neurons may reveal new therapeutic targets. It will also be important to explore whether similar patterns of synapse-specific susceptibility to dysregulation are observed in other epilepsy models or neurodevelopmental disorders.

## Acknowledgements

This work was supported by funding from the Brain Research Foundation, Seale Innovation Fund, Commonwealth Health Research Board, and NIH/NINDS (NS105804).

## Contributions

Conceptualization: M.S. and S.A.S.; Data curation: M.S., H.R., T.J.R, R.K., J.S., S.A.S.; Formal analysis: M.S., R.D., H.R., T.J.R., S.A.S.; Project administration, Supervision, Fund acquisition: S.A.S; Validation: M.S., R.D., R.K., S.A.S., Visualization: M.S., S.A.S.; Writing (original draft): M.S. and S.A.S; Writing (review and editing): M.S., R.D., H.R., T.J.R., H.R., R.K., J.S., S.A.S.

## Abbreviations

DS: Dravet Syndrome
CT: Corticothalamic
SUDEP: Sudden Unexpected Death
nRT: nucleus reticular of the thalamus
VPL: ventral posterolateral
VPM: ventral posteromedial thalamus.

