## Supplemental Figures S1-S8,Tables S1-S4 for "Input-and cell-type-specific developmental alterations to thalamic synapses in a Dravet syndrome mouse model"

**Corresponding Author:**

Sharon A. Swanger

**Contents:**

**Supplemental Figures S1 – S8**

**Supplemental Tables S1 – S4**

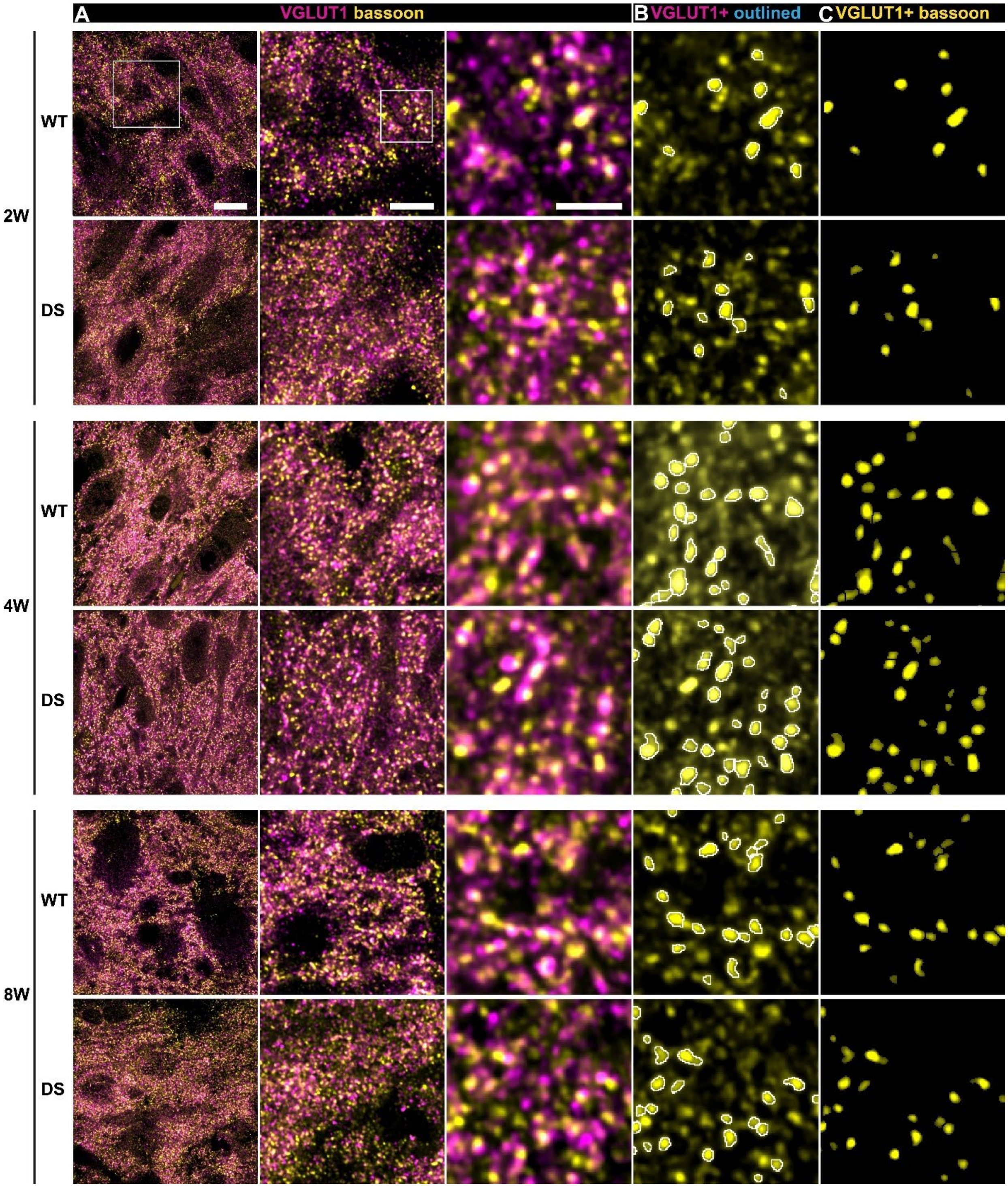


**Figure S1. Raw images for VGLUT1-positive synapse quantification in the VPL.**

**A.** 60x images with 4x SoRa of VGLUT1 and bassoon immunostained brain sections from WT and DS mice aged 2W, 4W, and 8W are shown. Regions of the full field image (left) were expanded to show synapse density (middle) and individual synapses (right). Expanded regions in subsequent rows are the same size. Scales: 10 µm, 5 µm, and 2 µm.

**B.** Image shows bassoon signal from the image in panel A with VGLUT1-positive puncta outlined (white).

**C.** Image shows only VGLUT1-positive bassoon puncta, which was overlaid with VGLUT1 in Figure 3.

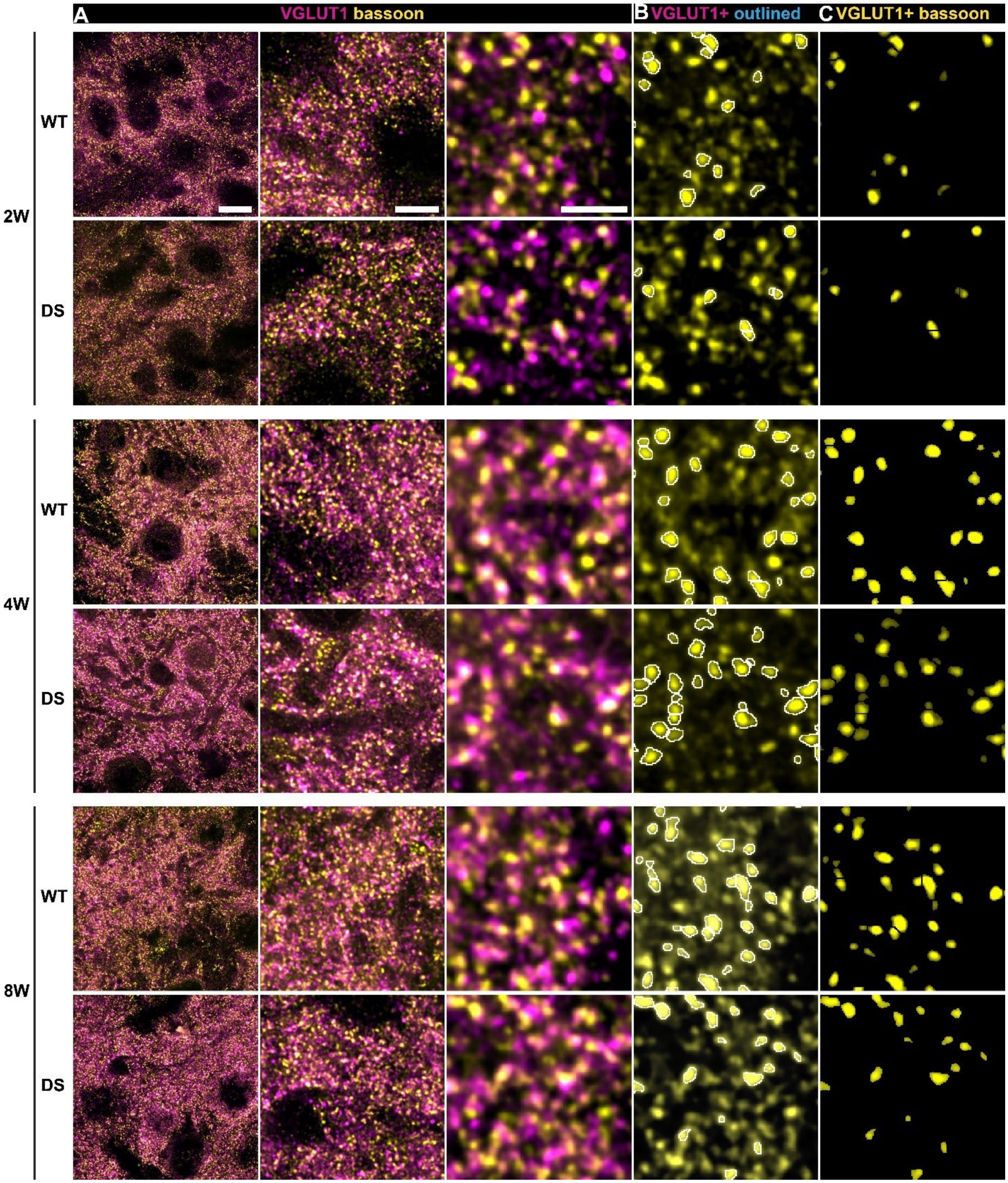


**Figure S2. Raw images for VGLUT1-positive synapse quantification in the VPM.**

**A.** 60x images with 4x SoRa of VGLUT1 and bassoon immunostained brain sections from WT and DS mice aged 2W, 4W, and 8W are shown. Regions of the full field image (left) were expanded to show synapse density (middle) and individual synapses (right). Expanded regions in subsequent rows are the same size. Scales: 10 µm, 5 µm, and 2 µm.

**B.** Image shows bassoon signal from the image in panel A with VGLUT1-positive puncta outlined (white).

**C.** Image shows only VGLUT1-positive bassoon puncta, which was overlaid with VGLUT1 in Figure 3.

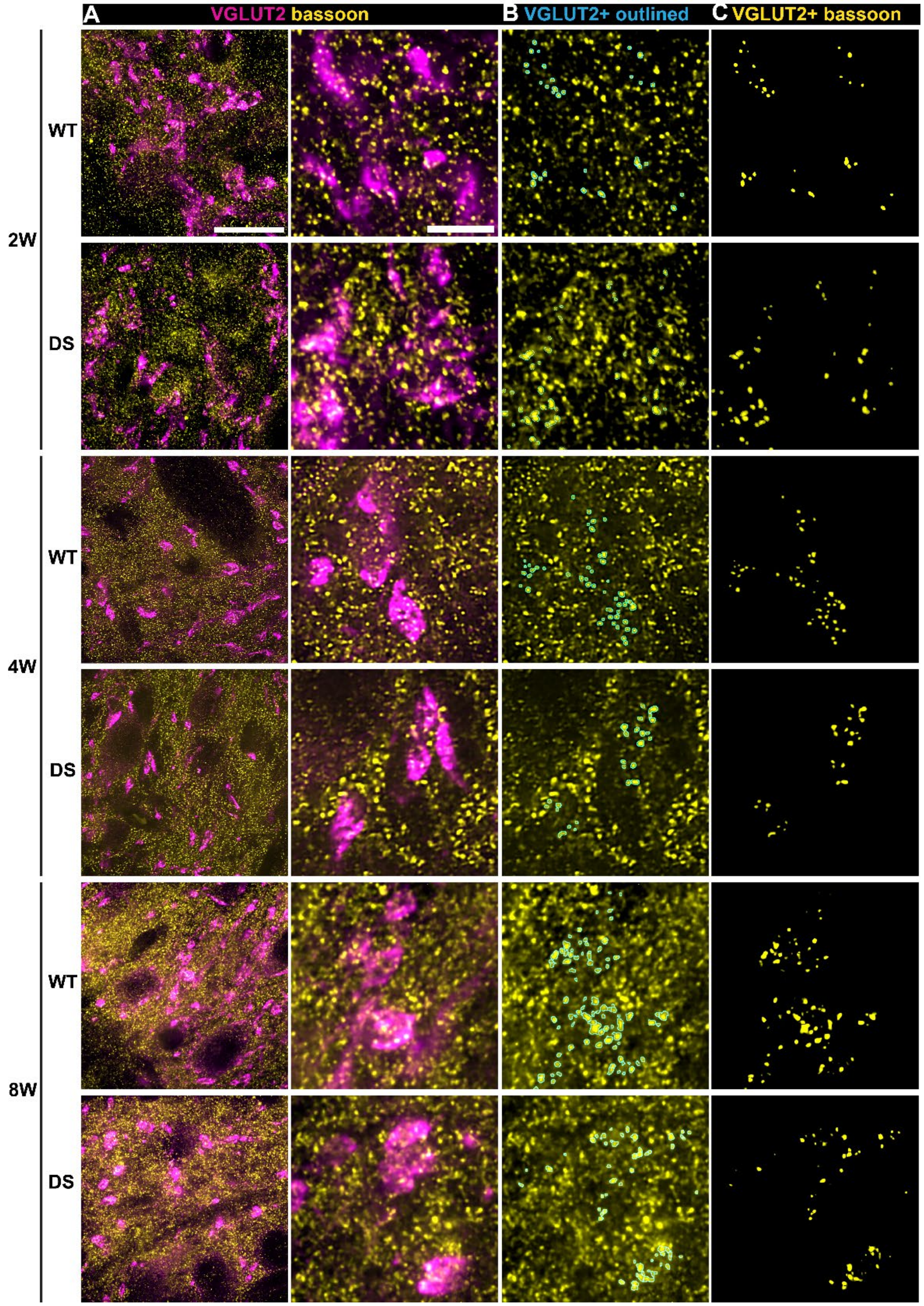


**Figure S3. Raw images for VGLUT2-positive synapse quantification in the VPL.**

**A.** 60x images with 4x SoRa of VGLUT2 and bassoon immunostained brain sections from WT and DS mice aged 2W, 4W, and 8W are shown. Regions of the full field image (left) were expanded to show axon terminals with multiple bassoon puncta (right). Expanded regions in subsequent rows are the same size. Scales: 10 µm and 2 µm.

**B.** Image shows bassoon signal from the image in panel A with VGLUT2-positive puncta outlined (cyan).

**C.** Image shows only VGLUT2-positive bassoon puncta, which was overlaid with VGLUT2 in Figure 4.

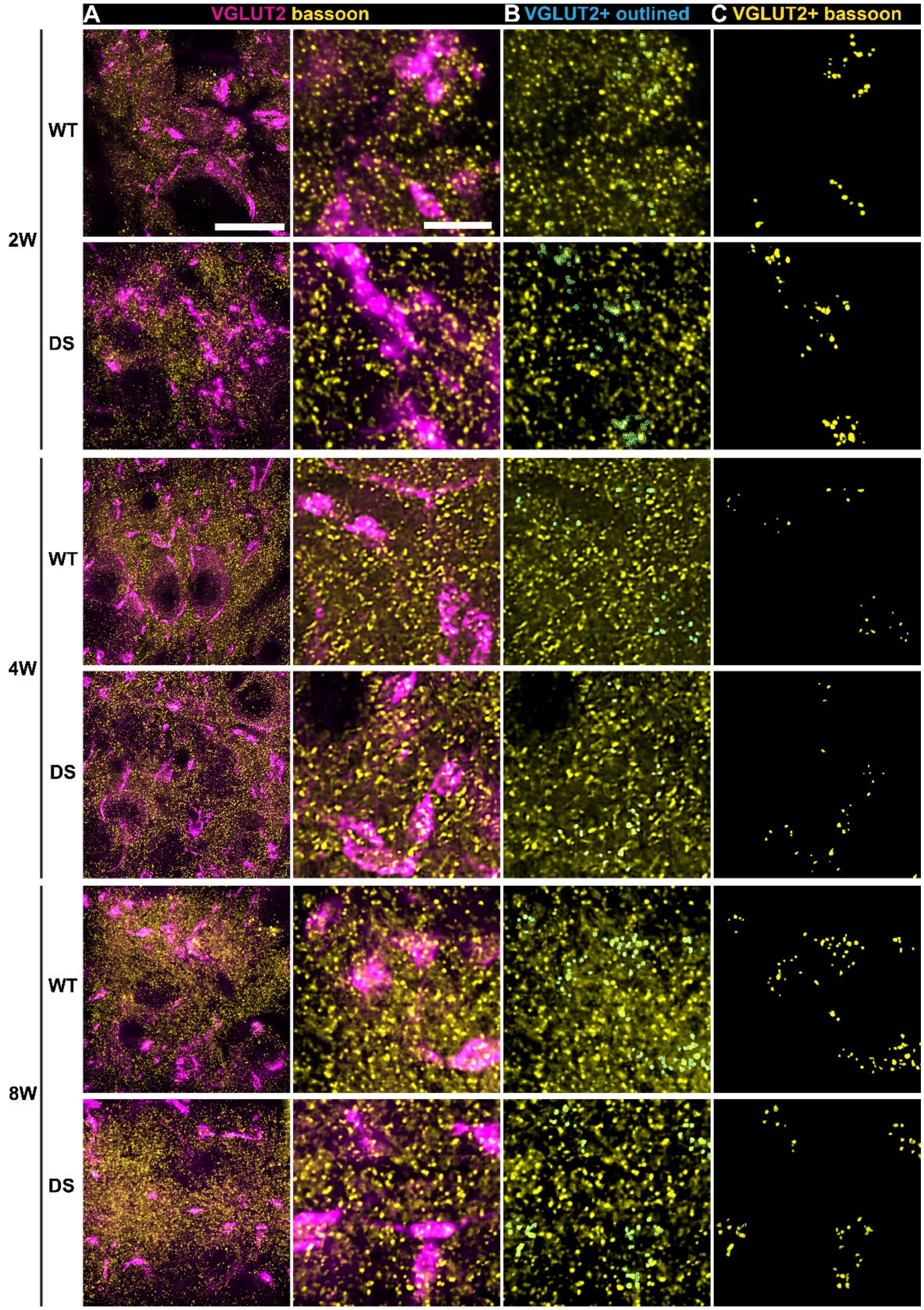


**Figure S4. Raw images for VGLUT2-positive synapse quantification in the VPM.**

**A.** 60x images with 4x SoRa of VGLUT2 and bassoon immunostained brain sections from WT and DS mice aged 2W, 4W, and 8W are shown. Regions of the full field image (left) were expanded to show axon terminal with multiple bassoon puncta (right). Expanded regions in subsequent rows are the same size. Scales: 10 µm and 2 µm.

**B.** Image shows bassoon signal from the image in panel A with VGLUT2-positive puncta outlined (cyan).

**C.** Image shows only VGLUT2-positive bassoon puncta, which was overlaid with VGLUT2 in Figure 4.

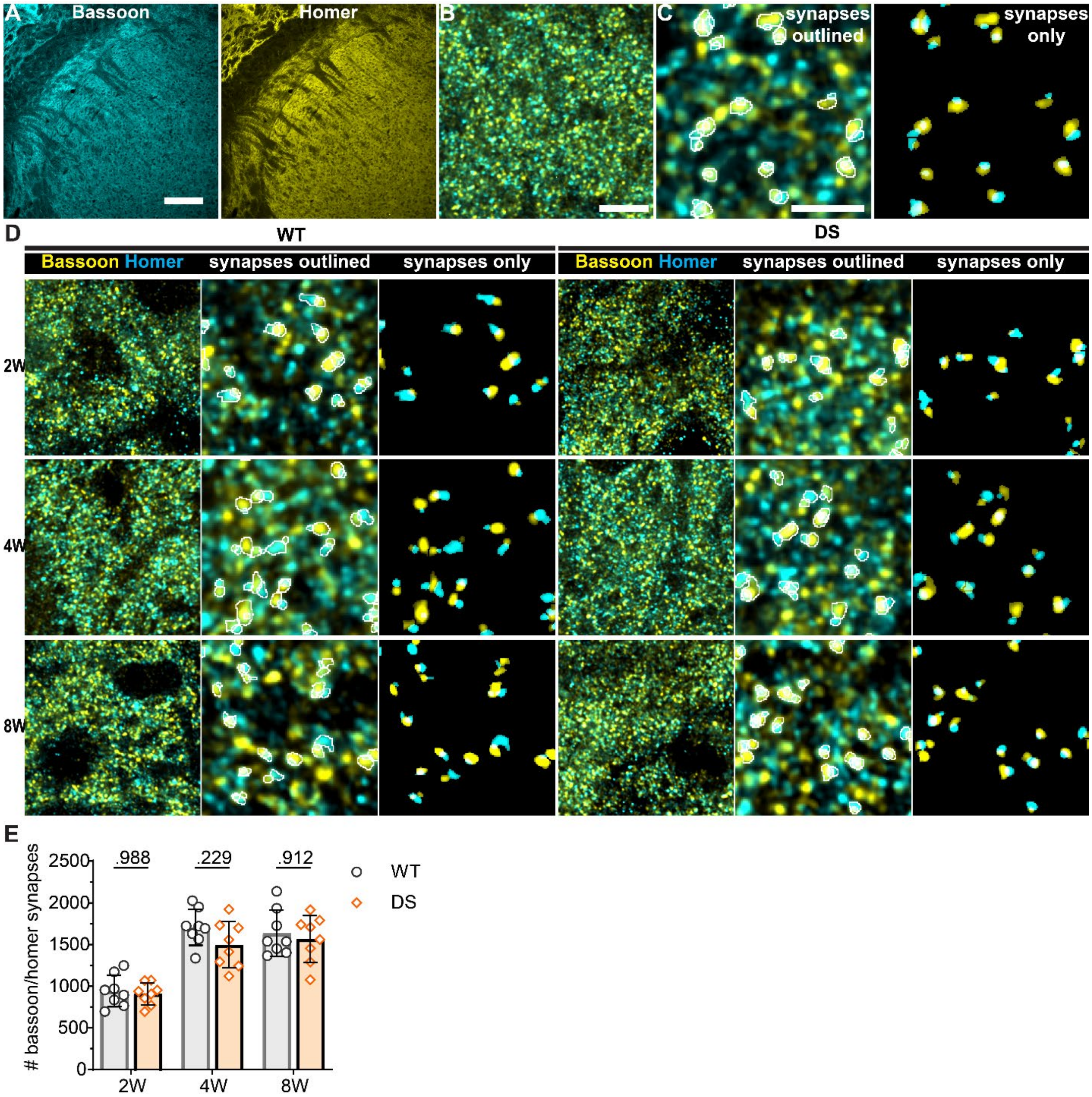


**Figure S5. Total VPL glutamatergic synapse number is similar between WT and DS mice.**

**A.** Representative 10x and **(B)** 60x (4x SoRa) images show homer and bassoon immunostaining of the thalamus. Scale: 200 µm and 10 µm.

**C.** Expanded region from the 60x (4x SoRa) image shows overlapping homer and bassoon puncta outlined (white, left) and only the puncta identified as synapses shown at right. Scale: 2 µm.

**D.** Representative ROIs from 60x images show homer and bassoon immunostained brain sections from WT and DS mice aged 2W (WT: n = 8, DS: n = 8), 4W (WT: n = 8, DS: n = 8), and 8W (WT: n = 8, DS: n = 8). Expanded images show overlapping pairs (synapses) outlined and synaptic puncta only (at right).

**E.** The number of synapses were averaged across three images from the VPL in each mouse. The group mean and SD were plotted with data points for all mice. Data were analyzed by one-way ANOVA with posthoc Sidak's tests between genotypes at each age (p-values shown in plots).

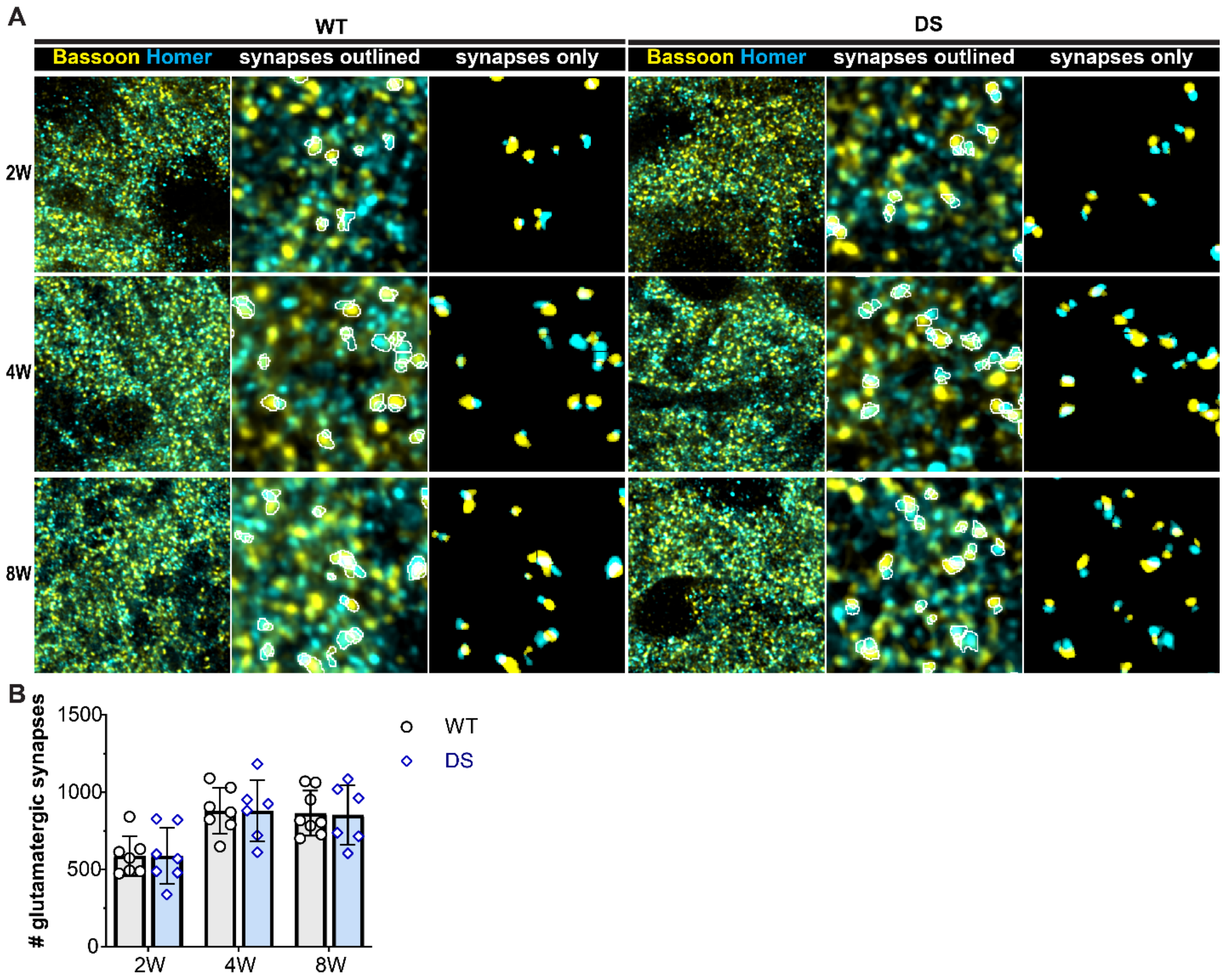


**Figure S6. Total VPM glutamatergic synapse number is similar between WT and DS mice.**

**A.** Representative 10x and **(B)** 60x (4x SoRa) images show homer and bassoon immunostaining of the thalamus. Scale: 200 µm and 10 µm.

**B.** Expanded region from the 60x (4x SoRa) image shows overlapping homer and bassoon puncta outlined (white, left) and only the puncta identified as synapses shown at right. Scale: 2 µm.

**C.** Representative ROIs from 60x images show homer and bassoon immunostained brain sections from WT and DS mice aged 2W (WT: n = 8, DS: n = 8), 4W (WT: n = 8, DS: n = 8), and 8W (WT: n = 8, DS: n = 8). Expanded images show overlapping pairs (synapses) outlined and synaptic puncta only (at right).

**D.** The number of synapses were averaged across three images from the VPM in each mouse. The group mean and SD were plotted with data points for all mice. Data were analyzed by one-way ANOVA with posthoc Sidak's tests between genotypes at each age (p-values shown in plots).

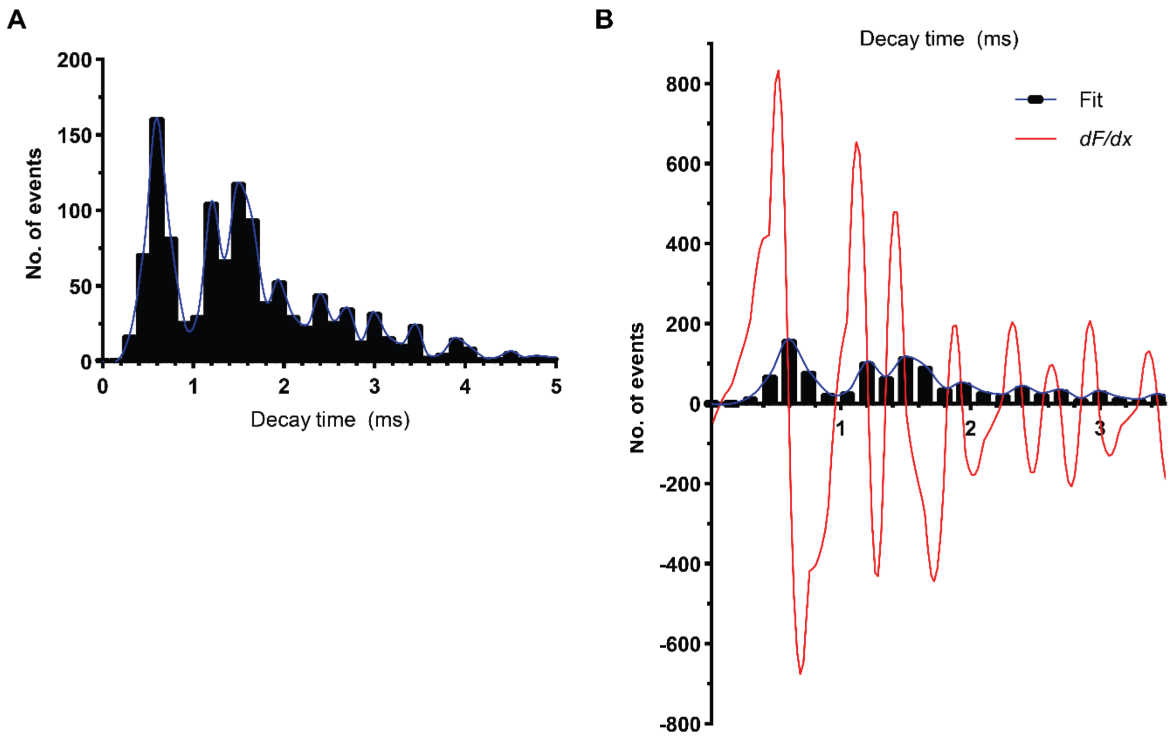


**Figure S7. Representative histograms for establishing Type 1 and Type 2 mEPSCs.**

**A.** Histogram shows total number of events vs decay time, with events binned across 0.2 ms. Data were interpoplated with a non-smoothing spline (blue line).

**B.** The derivative (*dF/dx,* red line) of the spline was plotted, and the decay time value where the *dF/dx* plot crosses zero between the two prominent histogram peaks was used at the cutoff value for Type 1 and Type 2 mEPSCs.

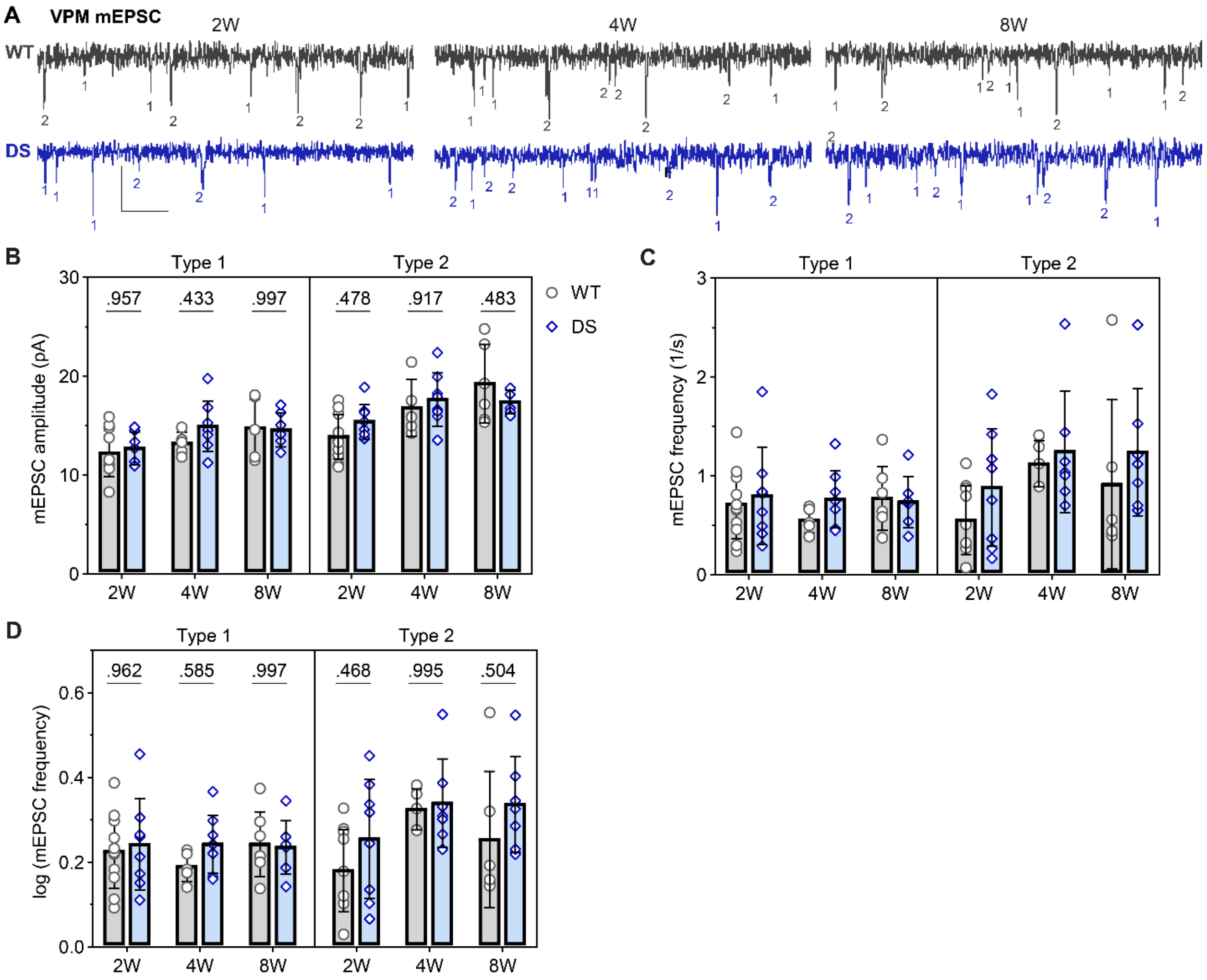


**Figure S8. The frequency and amplitude of VPM Type 1 and 2 mEPSC are similar between WT and DS mice.**

**A.** Representative voltage-clamp recordings from VPM neurons show mEPSCs in WT and DS mice aged P13 – P17 (WT: n = 11; DS: n = 7), P28 – P32 (WT: n = 7; DS: n = 8), P57 – P62 (WT: n = 7; DS: n = 7) with each mEPSC denoted as Type 1 (fast, putative sensory) or Type 2 (slow, putative CT) based on their decay kinetics.

**B.** Mean mEPSC amplitude were plotted for each mouse with the group mean ± *SD*. The data were analyzed by two-way ANOVA with pairwise Sidak's comparisons (p-values shown on plots) between WT and DS at each age.

**C.** The mean mEPSC frequency was plotted as raw data (geometric mean ± 95CI) and testing showed a non-normal distribution.

**D.** Transformed log10 values (mean ± 95CI) of the mean mESPC frequency were plotted and analyzed by two-way ANOVA with pairwise Sidak's comparisons (p-values shown on plots) between WT and DS at each age.

**Supplemental Table 1. Summary of electrophysiology normality and heteroscedasticity statistics.**

| | D'Agostino-Pearson test for Normality of residuals | | | | Spearman's test for heteroscedasticity | | | |
|---|---|---|---|---|---|---|---|---|
| | Raw Data | | Log transformed data | | Raw Data | | Log transformed data | |
| **EPSC Frequency** | K2 | p-value | K2 | p-value | R | p-value | R | p-value |
| VPL sEPSC | 18.00 | <.001 | 0.79 | 0.091 | 0.244 | 0.039 | 0.039 | 0.889 |
| VPM sEPSC | 14.23 | 0.001 | 3.33 | 0.190 | 0.539 | <.001 | 0.313 | 0.014 |
| VPL mEPSC | 19.37 | <.001 | 1.61 | 0.447 | 0.449 | <.001 | 0.043 | 0.382 |
| VPM mEPSC | 30.78 | <.001 | 0.96 | 0.081 | 0.169 | 0.128 | 0.540 | 0.358 |
| VPL mEPSC Type 1 | 23.43 | <.001 | 5.39 | 0.068 | 0.461 | <.001 | 0.247 | 0.051 |
| VPL mEPSC Type 2 | 17.57 | <.001 | 3.88 | 0.144 | 0.333 | 0.010 | 0.189 | 0.099 |
| VPM mEPSC Type 1 | 12.59 | 0.002 | 3.69 | 0.158 | 0.193 | 0.099 | 0.102 | 0.251 |
| VPM mEPSC Type 2 | 14.29 | 0.001 | 4.01 | 0.134 | -0.031 | 0.421 | -0.213 | 0.085 |
| **EPSC Amplitude** | K2 | p-value | | | R | p-value | | |
| VPL sEPSC | 4.29 | 0.117 | - | - | 0.088 | 0.262 | - | - |
| VPM sEPSC | 3.43 | 0.180 | - | - | 0.158 | 0.134 | - | - |
| VPL mEPSC | 1.68 | 0.431 | - | - | 0.030 | 0.409 | - | - |
| VPM mEPSC | 2.73 | 0.255 | - | - | 0.028 | 0.423 | - | - |
| VPL mEPSC Type 1 | 1.05 | 0.592 | - | - | 0.228 | 0.066 | - | - |
| VPL mEPSC Type 2 | 2.99 | 0.224 | - | - | 0.099 | 0.254 | - | - |
| VPM mEPSC Type 1 | 1.03 | 0.597 | - | - | -0.049 | 0.377 | - | - |
| VPM mEPSC Type 2 | 2.49 | 0.288 | - | - | 0.172 | 0.130 | - | - |
| **IPSC Frequency** | K2 | p-value | K2 | p-value | R | p-value | R | p-value |
| VPL sIPSC | 10.37 | 0.006 | 1.91 | 0.385 | 0.328 | 0.006 | -0.024 | 0.431 |
| VPM sIPSC | 7.95 | 0.019 | 1.09 | 0.579 | 0.322 | 0.009 | 0.034 | 0.404 |
| VPL mIPSC | 7.87 | 0.020 | 0.24 | 0.885 | 0.282 | 0.012 | -0.023 | 0.428 |
| VPM mIPSC | 7.48 | 0.024 | 1.02 | 0.601 | 0.462 | <.001 | 0.254 | 0.033 |
| **IPSC Amplitude** | K2 | p-value | | | K2 | p-value | | |
| VPL sIPSC | 5.14 | 0.077 | - | - | -0.114 | 0.202 | - | - |
| VPM sIPSC | 1.43 | 0.489 | - | - | -0.090 | 0.264 | - | - |
| VPL mIPSC | 2.81 | 0.245 | - | - | 0.105 | 0.199 | - | - |
| VPM mIPSC | 1.06 | 0.590 | - | - | 0.256 | 0.024 | - | - |

**Supplemental Table 2. Summary of electrophysiology data.**

| | Frequency | | | | | | | | | | | | Amplitude | | | | | |
|---|---|---|---|---|---|---|---|---|---|---|---|---|---|---|---|---|---|---|
| | Raw data: Geometric mean [95 CI] | | | | | | Transformed data: Mean [95 CI] | | | | | | Raw data: Mean (*SD*) | | | | | |
| | WT | | | DS | | | WT | | | DS | | | WT | | | DS | | |
| **VPL sEPSC** | GM | UL | LL | GM | UL | LL | M | UL | LL | M | UL | LL | M | SD | n | M | SD | n |
| 2W | 3.05 | 4.05 | 2.30 | 3.22 | 4.99 | 2.07 | 0.48 | 0.61 | 0.36 | 0.51 | 0.70 | 0.32 | 16.89 | 2.60 | 10 | 16.20 | 0.91 | 6 |
| 4W | 3.59 | 4.85 | 2.66 | 2.11 | 2.87 | 1.55 | 0.55 | 0.69 | 0.42 | 0.32 | 0.46 | 0.19 | 17.70 | 4.30 | 11 | 15.03 | 3.49 | 9 |
| 8W | 3.82 | 5.35 | 2.73 | 2.50 | 3.87 | 1.62 | 0.58 | 0.73 | 0.44 | 0.40 | 0.59 | 0.21 | 15.91 | 2.62 | 11 | 17.35 | 2.97 | 8 |
| **VPM sEPSC** | GM | UL | LL | GM | UL | LL | M | UL | LL | M | UL | LL | M | SD | n | M | SD | n |
| 2W | 1.55 | 1.93 | 1.24 | 1.57 | 2.21 | 1.11 | 0.41 | 0.47 | 0.35 | 0.42 | 0.51 | 0.32 | 14.21 | 1.45 | 9 | 14.03 | 2.32 | 8 |
| 4W | 2.30 | 3.40 | 1.56 | 2.23 | 2.87 | 1.73 | 0.53 | 0.65 | 0.41 | 0.51 | 0.59 | 0.44 | 17.35 | 4.15 | 9 | 16.03 | 2.86 | 11 |
| 8W | 2.13 | 2.97 | 1.52 | 2.28 | 3.59 | 1.45 | 0.50 | 0.60 | 0.40 | 0.53 | 0.66 | 0.39 | 16.72 | 2.66 | 7 | 17.30 | 2.55 | 7 |
| **VPL mEPSC** | GM | UL | LL | GM | UL | LL | M | UL | LL | M | UL | LL | M | SD | n | M | SD | n |
| 2W | 3.34 | 4.84 | 2.30 | 3.27 | 4.26 | 2.51 | 0.52 | 0.68 | 0.36 | 0.51 | 0.63 | 0.40 | 12.14 | 1.80 | 9 | 13.12 | 1.87 | 6 |
| 4W | 3.52 | 4.42 | 2.80 | 1.84 | 2.30 | 1.46 | 0.55 | 0.65 | 0.45 | 0.26 | 0.36 | 0.17 | 17.70 | 1.55 | 8 | 14.28 | 1.93 | 9 |
| 8W | 3.23 | 4.91 | 2.13 | 2.00 | 2.55 | 1.57 | 0.51 | 0.69 | 0.33 | 0.30 | 0.41 | 0.19 | 17.12 | 2.83 | 8 | 18.58 | 2.86 | 10 |
| **VPM mEPSC** | GM | UL | LL | GM | UL | LL | M | UL | LL | M | UL | LL | M | SD | n | M | SD | n |
| 2W | 1.60 | 2.23 | 1.15 | 1.90 | 2.80 | 1.29 | 0.20 | 0.35 | 0.06 | 0.28 | 0.45 | 0.11 | 12.50 | 2.25 | 11 | 14.07 | 1.80 | 8 |
| 4W | 1.83 | 2.65 | 1.27 | 1.82 | 2.43 | 1.36 | 0.26 | 0.42 | 0.10 | 0.26 | 0.39 | 0.13 | 13.93 | 2.60 | 7 | 13.10 | 1.98 | 8 |
| 8W | 2.12 | 3.07 | 1.47 | 2.43 | 3.53 | 1.67 | 0.33 | 0.49 | 0.17 | 0.39 | 0.55 | 0.22 | 13.90 | 2.87 | 7 | 12.88 | 2.00 | 9 |
| **VPL T1 mEPSC** | GM | UL | LL | GM | UL | LL | M | UL | LL | M | UL | LL | M | SD | n | M | SD | n |
| 2W | 1.16 | 1.59 | 0.84 | 1.02 | 1.28 | 0.82 | 0.34 | 0.41 | 0.26 | 0.31 | 0.36 | 0.26 | 12.36 | 1.33 | 6 | 11.71 | 1.37 | 9 |
| 4W | 1.94 | 2.57 | 1.47 | 0.76 | 0.93 | 0.62 | 0.47 | 0.56 | 0.39 | 0.25 | 0.29 | 0.21 | 16.07 | 2.53 | 8 | 13.45 | 1.52 | 8 |
| 8W | 1.61 | 2.87 | 0.91 | 1.06 | 1.44 | 0.78 | 0.43 | 0.58 | 0.29 | 0.32 | 0.39 | 0.25 | 14.77 | 2.13 | 7 | 16.40 | 2.13 | 8 |
| **VPM T1 mEPSC** | GM | UL | LL | GM | UL | LL | M | UL | LL | M | UL | LL | M | SD | n | M | SD | n |
| 2W | 0.63 | 0.91 | 0.44 | 0.69 | 1.11 | 0.42 | 0.23 | 0.28 | 0.17 | 0.24 | 0.33 | 0.15 | 12.20 | 2.34 | 11 | 12.68 | 1.65 | 7 |
| 4W | 0.54 | 0.73 | 0.40 | 0.72 | 0.98 | 0.53 | 0.19 | 0.23 | 0.15 | 0.24 | 0.30 | 0.19 | 13.20 | 1.15 | 7 | 14.93 | 2.54 | 8 |
| 8W | 0.72 | 1.05 | 0.49 | 0.70 | 0.97 | 0.50 | 0.24 | 0.31 | 0.17 | 0.24 | 0.29 | 0.18 | 14.77 | 2.85 | 7 | 14.56 | 1.71 | 7 |
| **VPL T2 mEPSC** | GM | UL | LL | GM | UL | LL | M | UL | LL | M | UL | LL | M | SD | n | M | SD | n |
| 2W | 1.81 | 2.69 | 1.21 | 1.80 | 2.44 | 1.33 | 1.36 | 1.63 | 1.10 | 1.37 | 1.59 | 1.14 | 14.15 | 2.47 | 6 | 12.85 | 1.56 | 9 |
| 4W | 1.68 | 2.19 | 1.29 | 1.19 | 1.66 | 0.85 | 1.31 | 1.48 | 1.14 | 1.11 | 1.29 | 0.92 | 17.11 | 2.40 | 8 | 16.10 | 2.12 | 7 |
| 8W | 1.52 | 2.45 | 0.95 | 1.41 | 1.72 | 1.16 | 1.28 | 1.56 | 1.01 | 1.20 | 1.32 | 1.07 | 16.89 | 2.90 | 9 | 18.70 | 2.49 | 8 |
| **VPM T2 mEPSC** | GM | UL | LL | GM | UL | LL | M | UL | LL | M | UL | LL | M | SD | n | M | SD | n |
| 2W | 0.43 | 0.78 | 0.24 | 0.67 | 1.38 | 0.32 | 0.18 | 0.25 | 0.11 | 0.25 | 0.37 | 0.14 | 13.87 | 2.26 | 11 | 15.40 | 1.78 | 8 |
| 4W | 1.10 | 1.43 | 0.85 | 1.15 | 1.69 | 0.78 | 0.32 | 0.38 | 0.26 | 0.34 | 0.44 | 0.24 | 16.81 | 2.87 | 5 | 17.64 | 2.69 | 8 |
| 8W | 0.70 | 1.52 | 0.32 | 1.12 | 1.73 | 0.73 | 0.25 | 0.42 | 0.08 | 0.34 | 0.44 | 0.23 | 19.24 | 3.98 | 6 | 17.41 | 1.18 | 7 |
| **VPL sIPSC** | GM | UL | LL | GM | UL | LL | M | UL | LL | M | UL | LL | M | SD | n | M | SD | n |
| 2W | 6.14 | 8.81 | 4.28 | 7.32 | 9.96 | 5.38 | 0.79 | 0.94 | 0.63 | 0.86 | 1.00 | 0.73 | 25.31 | 4.33 | 11 | 25.23 | 5.74 | 9 |
| 4W | 7.29 | 10.13 | 5.24 | 3.66 | 5.18 | 2.58 | 0.86 | 1.01 | 0.72 | 0.56 | 0.71 | 0.41 | 25.84 | 5.42 | 9 | 26.83 | 6.25 | 8 |
| 8W | 8.22 | 11.37 | 5.95 | 4.81 | 6.39 | 3.63 | 0.92 | 1.06 | 0.77 | 0.68 | 0.81 | 0.56 | 29.90 | 4.29 | 8 | 27.34 | 2.98 | 11 |
| **VPM sIPSC** | GM | UL | LL | GM | UL | LL | M | UL | LL | M | UL | LL | M | SD | n | M | SD | n |
| 2W | 8.69 | 10.98 | 6.88 | 6.55 | 10.28 | 4.18 | 0.94 | 1.04 | 0.84 | 0.82 | 1.01 | 0.62 | 27.11 | 6.58 | 10 | 22.13 | 2.57 | 9 |
| 4W | 9.19 | 13.31 | 6.35 | 4.14 | 5.99 | 2.86 | 0.96 | 1.12 | 0.80 | 0.62 | 0.78 | 0.46 | 28.47 | 6.24 | 11 | 26.07 | 4.56 | 9 |
| 8W | 7.45 | 11.35 | 4.88 | 6.57 | 9.24 | 4.67 | 0.87 | 1.06 | 0.69 | 0.82 | 0.97 | 0.67 | 28.78 | 1.58 | 7 | 28.68 | 7.74 | 6 |
| **VPL mIPSC** | GM | UL | LL | GM | UL | LL | M | UL | LL | M | UL | LL | M | SD | n | M | SD | n |
| 2W | 6.07 | 8.95 | 4.11 | 5.53 | 7.68 | 3.98 | 0.78 | 0.95 | 0.61 | 0.74 | 0.89 | 0.60 | 23.20 | 3.61 | 12 | 21.97 | 2.13 | 10 |
| 4W | 7.79 | 10.19 | 5.96 | 4.03 | 5.32 | 3.06 | 0.89 | 1.01 | 0.78 | 0.61 | 0.73 | 0.49 | 26.90 | 3.92 | 12 | 25.28 | 5.42 | 13 |
| 8W | 5.36 | 7.36 | 3.91 | 3.37 | 4.82 | 2.36 | 0.73 | 0.87 | 0.59 | 0.53 | 0.68 | 0.37 | 25.63 | 3.92 | 10 | 25.32 | 3.20 | 9 |
| **VPM mIPSC** | GM | UL | LL | GM | UL | LL | M | UL | LL | M | UL | LL | M | SD | n | M | SD | n |
| 2W | 4.72 | 6.54 | 3.40 | 6.25 | 9.33 | 4.18 | 0.67 | 0.82 | 0.53 | 0.80 | 0.97 | 0.62 | 21.29 | 3.18 | 11 | 21.46 | 2.83 | 10 |
| 4W | 4.85 | 6.53 | 3.60 | 6.24 | 10.34 | 3.76 | 0.69 | 0.81 | 0.56 | 0.80 | 1.01 | 0.58 | 22.77 | 3.71 | 11 | 25.14 | 4.81 | 10 |
| 8W | 3.71 | 4.40 | 3.13 | 4.84 | 6.89 | 3.40 | 0.57 | 0.64 | 0.50 | 0.69 | 0.84 | 0.53 | 27.33 | 4.06 | 8 | 28.25 | 5.51 | 10 |

**Supplemental Table 3. Electrophysiology data ANOVA tables.**

| | Frequency ANOVA table | | | Amplitude ANOVA table | | |
|---|---|---|---|---|---|---|
| **VPL sEPSC** | ANOVA table | F (DFn, DFd) | P value | ANOVA table | F (DFn, DFd) | P value |
| 2W | Interaction | F (2, 47) = 1.99 | P=.148 | Interaction | F (2, 49) = 2.064 | P=.138 |
| 4W | Age | F (2, 47) = 0.481 | P=.621 | Row Factor | F (2, 49) = 0.036 | P=.964 |
| 8W | Genotype | F (1, 47) = 5.98 | P=.018 | Column Factor | F (1, 49) = 0.556 | P=.459 |
| **VPM sEPSC** | ANOVA table | F (DFn, DFd) | P value | ANOVA table | F (DFn, DFd) | P value |
| 2W | Interaction | F (2, 44) = 0.114 | P=.893 | Interaction | F (2, 45) = 0.140 | P=.869 |
| 4W | Row Factor | F (2, 44) = 4.199 | P=.021 | Row Factor | F (2, 45) = 5.465 | P=.008 |
| 8W | Column Factor | F (1, 44) = 0.026 | P=.872 | Column Factor | F (1, 45) < 0.001 | P=.976 |
| **VPL mEPSC** | ANOVA table | F (DFn, DFd) | P value | ANOVA table | F (DFn, DFd) | P value |
| 2W | Interaction | F (2, 46) = 3.14 | P=.052 | Interaction | F (2, 45) = 6.354 | P=.004 |
| 4W | Row Factor | F (2, 46) = 2.80 | P=.071 | Row Factor | F (2, 45) = 22.15 | P<.001 |
| 8W | Column Factor | F (1, 46) = 12.8 | P<.001 | Column Factor | F (1, 45) = 0.264 | P=.610 |
| **VPM mEPSC** | ANOVA table | F (DFn, DFd) | P value | ANOVA table | F (DFn, DFd) | P value |
| 2W | Interaction | F (2, 41) = 0.205 | P=.794 | Interaction | F (2, 44) = 1.780 | P=.181 |
| 4W | Row Factor | F (2, 41) = 0.217 | P=.082 | Row Factor | F (2, 44) = 0.042 | P=.958 |
| 8W | Column Factor | F (1, 41) = 0.430 | P=.664 | Column Factor | F (1, 44) = 0.021 | P=.885 |
| **VPL T1 mEPSC** | ANOVA table | F (DFn, DFd) | P value | ANOVA table | F (DFn, DFd) | P value |
| 2W | Interaction | F (2, 39) = 4.356 | P=.020 | Interaction | F (2, 40) = 4.840 | P=.013 |
| 4W | Row Factor | F (2, 39) = 1.328 | P=.277 | Row Factor | F (2, 40) = 14.05 | P<.001 |
| 8W | Column Factor | F (1, 39) = 20.38 | P<.001 | Column Factor | F (1, 40) = 0.9523 | P=.335 |
| **VPM T1 mEPSC** | ANOVA table | F (DFn, DFd) | P value | ANOVA table | F (DFn, DFd) | P value |
| 2W | Interaction | F (2, 39) = 5.644 | P=.007 | Interaction | F (2, 38) = 0.641 | P=.533 |
| 4W | Row Factor | F (2, 39) = 2.764 | P=.075 | Row Factor | F (2, 38) = 4.245 | P=.022 |
| 8W | Column Factor | F (1, 39) = 31.01 | P<.001 | Column Factor | F (1, 38) = 0.972 | P=.331 |
| **VPL T2 mEPSC** | ANOVA table | F (DFn, DFd) | P value | ANOVA table | F (DFn, DFd) | P value |
| 2W | Interaction | F (2, 42) = 0.626 | P=.540 | Interaction | F (2, 41) = 2.127 | P=.132 |
| 4W | Row Factor | F (2, 42) = 1.568 | P=.221 | Row Factor | F (2, 41) = 13.48 | P<.001 |
| 8W | Column Factor | F (1, 42) = 1.698 | P=.200 | Column Factor | F (1, 41) = 0.057 | P=.813 |
| **VPM T2 mEPSC** | ANOVA table | F (DFn, DFd) | P value | ANOVA table | F (DFn, DFd) | P value |
| 2W | Interaction | F (2, 37) = 0.315 | P=.732 | Interaction | F (2, 39) = 1.796 | P=.179 |
| 4W | Row Factor | F (2, 37) = 3.708 | P=.034 | Row Factor | F (2, 39) = 9.070 | P<.001 |
| 8W | Column Factor | F (1, 37) = 2.468 | P=.125 | Column Factor | F (1, 39) = 0.055 | P=.817 |
| **VPL sIPSC** | ANOVA table | F (DFn, DFd) | P value | ANOVA table | F (DFn, DFd) | P value |
| 2W | Interaction | F (2, 51) = 5.154 | P=.009 | Interaction | F (2, 50) = 0.636 | P=.534 |
| 4W | Row Factor | F (2, 51) = 1.746 | P=.185 | Row Factor | F (2, 50) = 2.36 | P=.105 |
| 8W | Column Factor | F (1, 51) = 8.692 | P=.005 | Column Factor | F (1, 50) = 0.179 | P=.674 |
| **VPM sIPSC** | ANOVA table | F (DFn, DFd) | P value | ANOVA table | F (DFn, DFd) | P value |
| 2W | Interaction | F (2, 47) = 2.459 | P=.096 | Interaction | F (2, 46) = 0.8210 | P=.446 |
| 4W | Row Factor | F (2, 47) = 0.8082 | P=.452 | Row Factor | F (2, 46) = 2.474 | P=.095 |
| 8W | Column Factor | F (1, 47) = 9.487 | P=.003 | Column Factor | F (1, 46) = 2.707 | P=.107 |
| **VPL mIPSC** | ANOVA table | F (DFn, DFd) | P value | ANOVA table | F (DFn, DFd) | P value |
| 2W | Interaction | F (2, 58) = 2.017 | P=.142 | Interaction | F (2, 60) = 0.154 | P=.858 |
| 4W | Row Factor | F (2, 58) = 2.523 | P=.089 | Row Factor | F (2, 60) = 5.093 | P=.009 |
| 8W | Column Factor | F (1, 58) = 11.45 | P=.001 | Column Factor | F (1, 60) = 1.171 | P=.284 |
| **VPM mIPSC** | ANOVA table | F (DFn, DFd) | P value | ANOVA table | F (DFn, DFd) | P value |
| 2W | Interaction | F (2, 47) = 0.008 | P=.992 | Interaction | F (2, 54) = 0.390 | P=.679 |
| 4W | Row Factor | F (2, 47) = 1.750 | P=.185 | Row Factor | F (2, 54) = 11.8 | P<.001 |
| 8W | Column Factor | F (1, 47) = 5.310 | P=.026 | Column Factor | F (1, 54) = 1.17 | P=.284 |

**Supplemental Table 4. Summary of imaging data and statistics.**

| | Raw data: Mean (SD) | | | | | | D'Agostino-Pearson test | | Spearman's test | | ANOVA table | | |
|---|---|---|---|---|---|---|---|---|---|---|---|---|---|
| | WT | | | DS | | | WT | DS | WT | DS | | | |
| **VPL # total** | M | SD | n | M | SD | n | K2 | p-value | R | p-value | ANOVA table | F (DFn, DFd) | p-value |
| 2W | 943 | 189 | 8 | 908 | 133 | 8 | | | | | Interaction | (2, 42) = 0.6[illegible] | P=.547 |
| 4W | 1706 | 216 | 8 | 1498 | 277 | 8 | 0.48 | 0.788 | 0.17 | 0.119 | Age | (2, 42) = 44.[illegible] | P<.001 |
| 8W | 1637 | 275 | 8 | 1568 | 280 | 8 | | | | | Genotype | (1, 42) = 2.3 | P=.131 |
| **VPM # total** | M | SD | n | M | SD | n | K2 | p-value | R | p-value | ANOVA table | F (DFn, DFd) | p-value |
| 2W | 995 | 172 | 8 | 923 | 154 | 8 | | | | | Interaction | (2, 42) = 0.09 | P=.914 |
| 4W | 1535 | 199 | 8 | 1417 | 210 | 8 | 5.75 | 0.056 | 0.18 | 0.110 | Row Factor | (2, 42) = 28.[illegible] | P<.001 |
| 8W | 1396 | 258 | 8 | 1267 | 201 | 8 | | | | | Column Factor | (1, 42) = 3.3 | P=.074 |
| **VPL # VGLUT1** | M | SD | n | M | SD | n | K2 | p-value | R | p-value | ANOVA table | F (DFn, DFd) | p-value |
| 2W | 907 | 217 | 7 | 1042 | 197 | 7 | | | | | Interaction | (2, 40) = 1.35 | P=.270 |
| 4W | 1772 | 311 | 8 | 1631 | 312 | 8 | 0.43 | 0.806 | 0.17 | 0.135 | Row Factor | (2, 40) = 33.5 | P<.001 |
| 8W | 1637 | 170 | 8 | 1508 | 281 | 8 | | | | | Column Factor | (1, 40) = 0.35 | P=.556 |
| **VPM # VGLUT1** | M | SD | n | M | SD | n | K2 | p-value | R | p-value | ANOVA table | F (DFn, DFd) | p-value |
| 2W | 785 | 176 | 7 | 1069 | 145 | 7 | | | | | Interaction | (2, 39) = 1.9 | P=.163 |
| 4W | 1346 | 286 | 7 | 1327 | 283 | 8 | 0.32 | 0.852 | -0.10 | 0.257 | Row Factor | (2, 39) = 22.[illegible] | P<.001 |
| 8W | 1335 | 187 | 8 | 1479 | 119 | 8 | | | | | Column Factor | (1, 39) = 4.8 | P=.035 |
| **VPL # VGLUT2** | M | SD | n | M | SD | n | K2 | p-value | R | p-value | ANOVA table | F (DFn, DFd) | p-value |
| 2W | 148 | 30 | 7 | 135 | 30 | 7 | | | | | Interaction | (2, 40) = 3.3 | P=.045 |
| 4W | 187 | 34 | 8 | 123 | 25 | 8 | 2.79 | 0.248 | 0.19 | 0.099 | Row Factor | (2, 40) = 0.7[illegible] | P=.500 |
| 8W | 191 | 46 | 8 | 117 | 35 | 8 | | | | | Column Factor | (1, 40) = 25.[illegible] | P<.001 |
| **VPM # VGLUT2** | M | SD | n | M | SD | n | K2 | p-value | R | p-value | ANOVA table | F (DFn, DFd) | p-value |
| 2W | 97 | 30 | 7 | 113 | 27 | 7 | | | | | Interaction | (2, 39) = 0.7[illegible] | P=.485 |
| 4W | 108 | 16 | 8 | 101 | 17 | 8 | 0.25 | 0.882 | 0.08 | 0.300 | Row Factor | (2, 39) = 0.18 | P=.833 |
| 8W | 107 | 29 | 7 | 112 | 27 | 8 | | | | | Column Factor | (1, 39) = 0.4[illegible] | P=.498 |
